# Highly sensitive in situ proteomics with cleavable fluorescent tyramide reveals human neuronal heterogeneity

**DOI:** 10.1101/539106

**Authors:** Renjie Liao, Manas Mondal, Christopher D. Nazaroff, Diego Mastroeni, Paul D. Coleman, Joshua LaBaer, Jia Guo

## Abstract

The ability to comprehensively profile proteins in intact tissues in situ is crucial for our understanding of health and disease. However, the existing methods suffer from low sensitivity and limited sample throughput. To address these issues, here we present a highly sensitive and multiplexed in situ protein analysis approach using cleavable fluorescent tyramide and off-the-shelf antibodies. Compared with the current methods, this approach enhances the detection sensitivity and reduces the imaging time by 1-2 orders of magnitude, and can potentially detect hundreds of proteins in intact tissues at the optical resolution. Applying this approach, we studied protein expression heterogeneity in genetically identical cells, and performed expression correlation analysis to identify coregulated proteins. We also profiled >6000 neurons in human formalin-fixed paraffin-embedded (FFPE) hippocampus. By partitioning these neurons into varied cell clusters based on their protein expression profiles, we observed different subregions of the hippocampus consist of neurons from distinct clusters.

## Introduction

Comprehensive protein profiling in individual cells of intact tissues in situ holds great promise to unlock major mysteries in neuroscience, cancer and stem cell biology^1^, since it can reveal the spatial organization, gene expression regulation, and interactions of the diverse cell types in complex multicellular organisms. Mass spectrometry^2^ and microarray technologies^3^ have been widely used for proteomics analysis. However, as these approaches are carried out on extracted and purified proteins from populations of cells, they lose the protein location information and conceal the single-cell expression variations in the sample. Fluorescence microscopy is a powerful tool to study protein expressions in individual cells in their native spatial contexts. However, due to the spectral overlap of organic fluorophores^4–6^ and fluorescent proteins^7,8^, conventional protein imaging technologies only allow a handful of proteins to be detected in one specimen.

To enable multiplexed single-cell protein analysis, a number of methods have been explored recently, such as mass cytometry^9^, single cell barcode chips^10–12^, and DNA-antibody barcoded arrays^13^. Nonetheless, as protein spatial complexity are masked in these approaches, they can not be applied to profile proteins in intact tissues in situ^14^. To address this issue, cyclical immunofluorescence^15–23^ and mass cytometry imaging^24,25^ have been developed. However, with the detection tags directly conjugated to antibodies, these existing methods have low detection sensitivity. This limitation hinders their applications to study proteins with low expression levels. Additionally, the low sensitivity of the current methods also limit their ability to profile proteins in highly autofluorescent tissues, such as formalin-fixed paraffin-embedded (FFPE) tissues^26^, which are the most common type of preserved clinical samples^27^. Moreover, the existing methods have limited sample throughput, as they require pixel-by-pixel sample analysis^24,25^ or high magnification objectives and long exposure time to detect the protein targets^15–23^.

Here, we report a highly sensitive and multiplexed in situ protein analysis approach with cleavable fluorescent tyramide (CFT). In this approach, target proteins are sensitively detected by a signal amplification method using off-the-shelf horseradish peroxidase (HRP) conjugated antibodies and CFT. Upon continuous cycles of target staining, fluorescence imaging, fluorophore cleavage and HRP deactivation, this approach has the potential to quantify hundreds of different proteins in individual cells of intact tissues at the optical resolution. To demonstrate the feasibility of this approach, we designed and synthesized CFT. We showed that the detection sensitivity and sample throughput of our approach are orders of magnitude higher than those of the existing methods. We also demonstrated that tris(2-carboxyethyl)phosphine (TCEP) can efficiently cleave the fluorophores and simultaneously deactivate HRP, while maintaining protein targets antigenicity. We validated our approach in HeLa cells and showed excellent agreement with conventional immunohistochemistry (IHC) results. Using this approach, we studied protein expression heterogeneity in a population of genetically identical cells, and performed the protein expression correlation analysis to identify coregulated proteins. We also applied this approach to investigate the neuronal heterogeneity in the human hippocampus, and demonstrated that distinct subregions of the hippocampus are composed of varied neuron clusters.

## Results

### Platform design

As shown in Fig. 1A, this protein profiling technology consists of three major steps in each analysis cycle. First, protein targets are recognized by HRP conjugated antibodies. And HRP catalyzes the coupling reaction between CFT and the tyrosine residues on the endogenous proteins in close proximity. In the second step, fluorescence images are captured to generate quantitative protein expression profiles. Finally, the fluorophores attached to tyramide are chemically cleaved and simultaneously HRP is deactivated, which allows the initiation of the next analysis cycle. Through reiterative cycles of target staining, fluorescence imaging, fluorophore cleavage and HRP deactivation, a large number of different proteins with a wide range of expression levels can be quantified in single cells of intact tissues in situ.

**Fig. 1.**
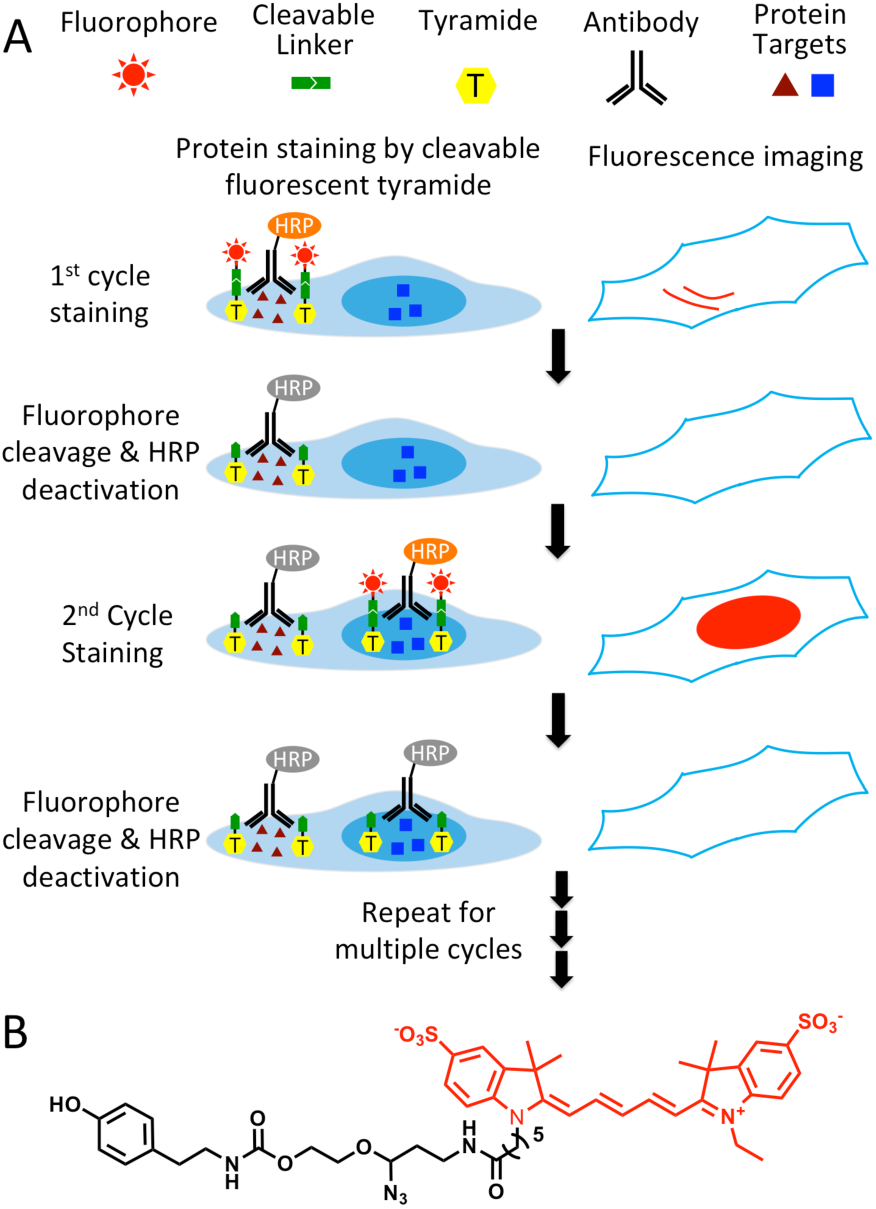
A) Highly sensitive and multiplexed in situ protein profiling with cleavable fluorescent tyramide. Protein targets are stained with HRP conjugated antibodies and cleavable fluorescent tyramide. After imaging, the fluorophores are chemically cleaved and simultaneously the HRP is deactivated. Through cycles of target staining, fluorescence imaging, fluorophore cleavage and HRP deactivation, comprehensive protein profiling can be achieved in single cells in situ. B) Structure of cleavable fluorescent tyramide, tyramide-N3-Cy5.

### Design and synthesis of CFT

To enable fluorescence signal removal after protein staining, we designed CFT by tethering fluorophores to tyramide through a chemically cleavable linker. A critical requirement for the success of this technology is to efficiently cleave the fluorophores in the cellular environment while maintaining the protein antigenicity. Additionally, it is preferred that the linker has a small size, so that HRP can still recognize CFT as a good substrate and the diffusion of short-lived tyramide radical^28^ is not compromised. Recently, our laboratory has developed an azide-based cleavable linker^21^, which satisfies all of those requirements. Thus, we incorporated that linker into CFT. Most of tissues exhibit prominent autofluorescence from the green and yellow emission channels, while only minimal autofluorescence is detected in the red emission channel^29^. To avoid the significant green and yellow autofluorescence background, in the current study we selected Cy5 as the fluorophore for CFT.

Synthesis of CFT (tyramide-N_3_-Cy5) (Fig. 1B) was carried out by coupling of tyramine and the Cy5 conjugated cleavable linker (Scheme S1). After purified by high performance liquid chromatography (HPLC), the prepared CFT was characterized by mass spectrometry and nuclear magnetic resonance (NMR) spectroscopy. The detailed synthesis and characterization of CFT is described in the supporting information.

### Significantly enhanced detection sensitivity

We next assessed the detection sensitivity of our approach by comparing it with direct and indirect immunofluorescence, which have similar sensitivity to the current multiplexed in situ protein profiling approaches^14^. Applying conventional immunofluorescence methods, we stained protein Ki67 in HeLa cells with Cy5 labeled primary antibodies (Fig. 2A), and unlabeled primary antibodies together with Cy5 labeled secondary antibodies (Fig. 2B). Using our approach, protein Ki67 was stained with unlabeled primary antibodies and HRP conjugated secondary antibodies along with tyramide-N_3_-Cy5 (Fig. 2C). With primary antibodies of the same concentration, our method is ∼88 and ∼35 times more sensitive than direct and indirect immunofluorescence, respectively (Fig. 2D). Additionally, the staining resolution of the three methods closely resembles each other (Fig. 2A-C). These results suggest that HRP can still recognize CFT as a good substrate and the incorporation of the cleavable linker into CFT does not interfere with the diffusion of the CFT radical. More importantly, the extremely high sensitivity of our approach enables the quantitative in situ analysis of low-abundance proteins, which could be undetectable by the reported methods. Moreover, by reducing the imaging time by 1-2 orders of magnitude, our method allows a large number of individual cells to be profiled in a short time, which leads to the dramatically improved sample throughput and minimized assay time.

**Fig. 2.**
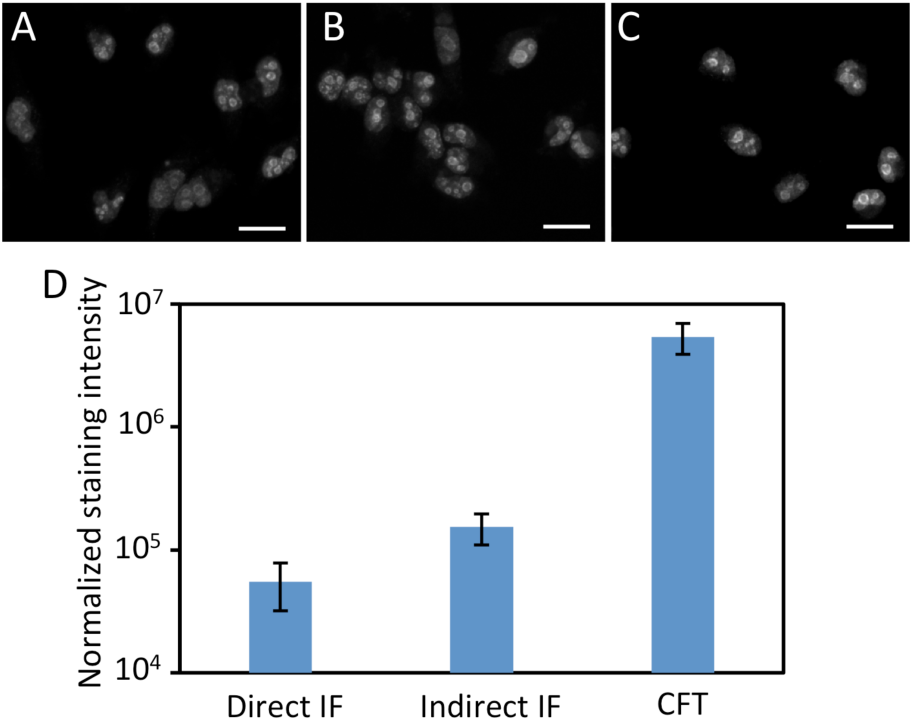
Protein Ki67 in HeLa cells are stained by (A) direct immunofluorescence (IF), (B) indirect IF, and (C) cleavable fluorescent tyramide (CFT). The images in (A), (B) and (C) are captured with the exposure time of 1 second, 300 millisecond, and 15 millisecond, respectively. (D) Normalized staining intensities of 30 different positions in (A), (B) and (C). The y-axis in (D) is on a logarithmic scale. Scale bars, 25 μm.

### Efficient fluorophore cleavage without loss of protein antigeneity

We next explored whether the fluorophores can be efficiently cleaved while maintaining the protein antigenicity. To search for this ideal cleavage condition, we stained protein Ki67 in HeLa cells using HRP conjugated antibodies and tyramide-N_3_-Cy5, and evaluated the fluorophore cleavage efficiencies at different temperature. After incubating with tris(2-carboxyethyl)phosphine (TCEP) at 37°C, 50°C and 65°C for 30 minutes, over 85%, 90% and 95% of the staining signals were removed, respectively (Supplementary Fig. 1). To test whether the protein antigenicity remains at those varied cleavage temperature, we incubated HeLa cells with TCEP at 37°C, 50°C and 65°C for 24 hours, and subsequently stained protein Ki67 with tyramide-N_3_-Cy5. We also labeled protein Ki67 without any pre-treatment as controls. The cells with the TCEP incubation at 37°C and 50°C have similar staining intensities to the control cells; while the cells pretreated at 65°C only have about half of the staining intensities compared to the control cells (Supplementary Fig. 2). We then studied the fluorophore cleavage kinetics at 50°C by incubating the stained cells with TCEP for 5, 15, 30 and 60 minutes. Among these conditions, 30 minutes is the minimum cleavage time required to achieve the maximum cleavage efficiency (Supplementary Fig. 3). These results indicate that the fluorescence signals generated by staining with CFT can be efficiently removed by the TCEP treatment at 50°C for 30 minutes, and this condition preserves the protein antigenicity.

### Simultaneous fluorophore cleavage and HRP activation

Another critical requirement for the success of this approach is that HRP needs to be deactivated at the end of each analysis cycle, so that it will not generate false positive signals in the next cycle. To explore whether TCEP can deactivate HRP and cleave fluorophores simultaneously, we stained proteins ILF3 (Fig. 3A), HMGB1, HDAC2, TDP43, PABPN1, hnRNP A1, Nucleolin, H4K16ac, hnRNP K and Nucleophosmin (Supplementary Fig. 4) in HeLa cells using HRP conjugated antibodies and tyramide-N_3_-Cy5. After TCEP incubation at 50°C for 30 minutes, the fluorescence signals were efficiently removed, yielding the on/off ratios of over 10:1 (Fig. 3B,D, Supplementary Fig. 4). We then incubated the cells with tyramide-N_3_-Cy5 again. For all the proteins under study, no further fluorescence signal increases were observed (Fig. 3C,D, Supplementary Fig. 4). These results confirm that the protein staining signals generated by CFT can be efficiently erased by TCEP, and also indicate that TCEP can deactivate HRP simultaneously.

**Fig. 3.**
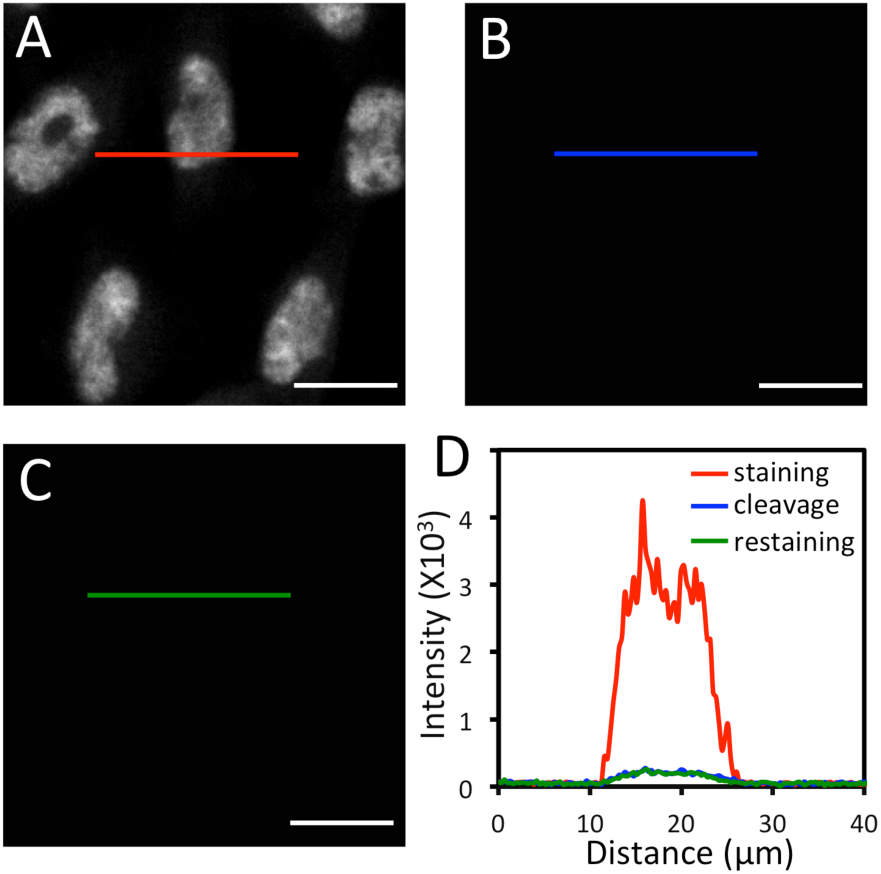
A) Protein ILF3 in HeLa cells is stained with HRP conjugated antibodies and tyramide-N3-Cy5. B) Cy5 is cleaved by TCEP. C) Cells are incubated with tyramide-N3-Cy5, again. D) Fluorescence intensity profile corresponding to the red, blue and green line positions in (A), (B) and (C). Scale bars, 20 μm.

### Multiplexed in situ protein profiling in HeLa cells

To demonstrate the feasibility of applying CFT for multiplexed protein analysis, we labeled 10 distinct proteins in individual HeLa cells in situ. Through reiterative staining cycles, proteins HMGB1, HDAC2, TDP43, PABPN1, hnRNP A1, Nucleolin, H4K16ac, hnRNP K, ILF3 and Nucleophosmin were unambiguously detected with the HRP conjugated antibodies and tyramide-N_3_-Cy5 in the same set of cells (Fig. 4). We also stained these 10 protein targets in 10 different sets of cells by conventional tyramide signal amplification (TSA) assays using Cy5 labeled tyramide (Supplementary Fig. 5). The protein distribution patterns obtained by the two methods are consistent with each other. We also compared the mean protein abundances per cell measured by our CFT-based approach and conventional immunofluorescence with TSA. For all the 10 proteins with varied expression levels, the results obtained using the two methods closely resemble each other (Fig. 5A). Comparison of the two sets of results yields an R^2^ value of 0.99 with a slope of 1.13 (Fig. 5B). These results indicate that our approach allows quantitative and multiplexed protein profiling in individual cells in situ.

**Fig. 4.**
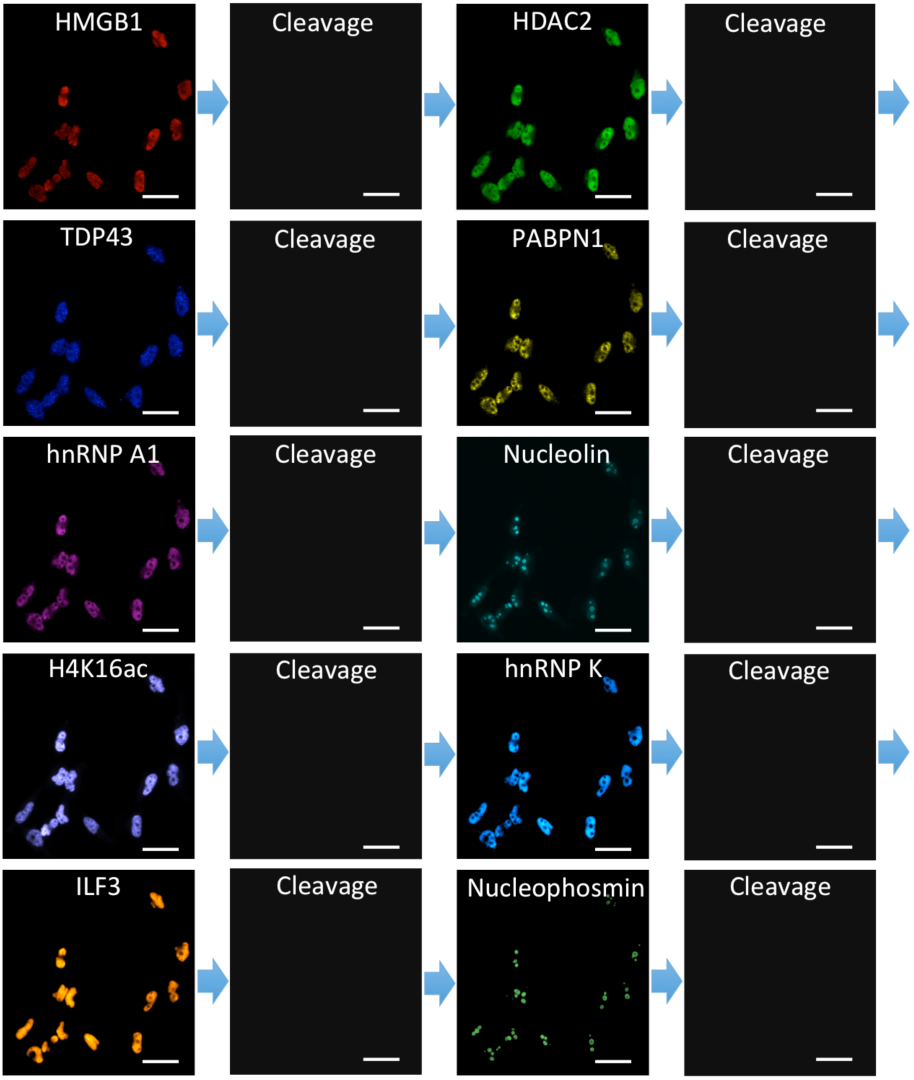
10 different proteins are stained sequentially with the corresponding HRP conjugated antibodies and tyramide-N3-Cy5 in the same set of HeLa cells. Scale bars, 40 μm.

**Fig. 5.**
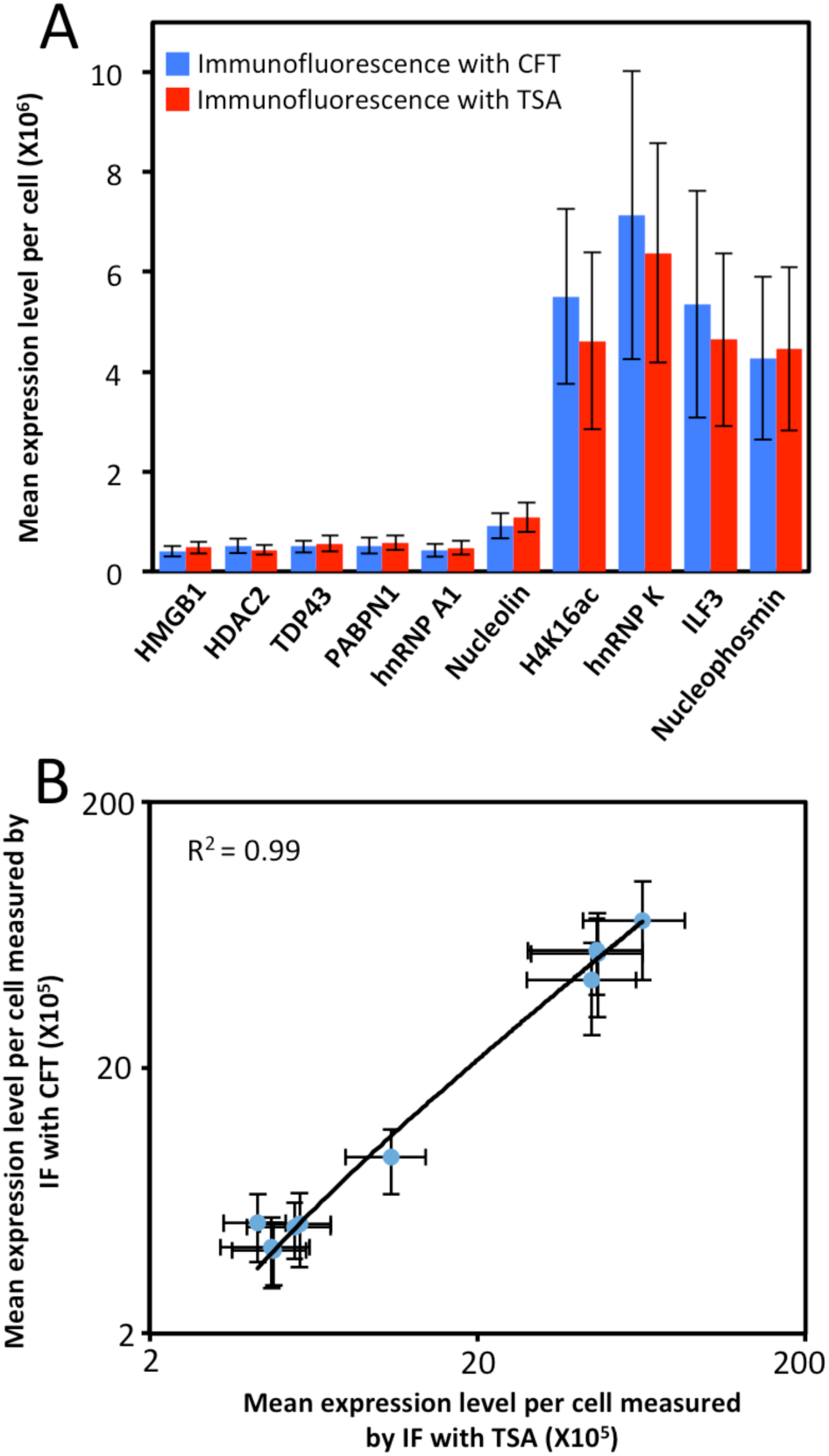
(A) Mean expression level per cell (n = 200 cells) of 10 different proteins measured by immunofluorescence (IF) with cleavable fluorescent tyramide (CFT) and conventional immunofluorescence with tyramide signal amplification (TSA). (B) Comparison of the results obtained by immunofluorescence with CFT and TSA yields R2 = 0.99 with a slope of 1.13. The x and y axes in (B) are on a logarithmic scale.

### Protein expression heterogeneity and correlation

As shown in many experiments, genetically identical cells can exhibit significant gene expression variations among individual cells^30–36^. To explore such cell-to-cell protein expression heterogeneity in HeLa cells, we analyzed the distribution of the single-cell protein expression levels. As shown in Fig. 6A, the single-cell protein abundances are distributed in a wide range. This significant expression heterogeneity results in the relatively large error bars in Fig. 5. For all the ten measured proteins, the square of the expression standard deviation is much higher than the mean expression levels (Fig. 6A). These results suggest that these proteins are generated in translational bursts, rather than at a constant rate^37^.

**Fig. 6.**
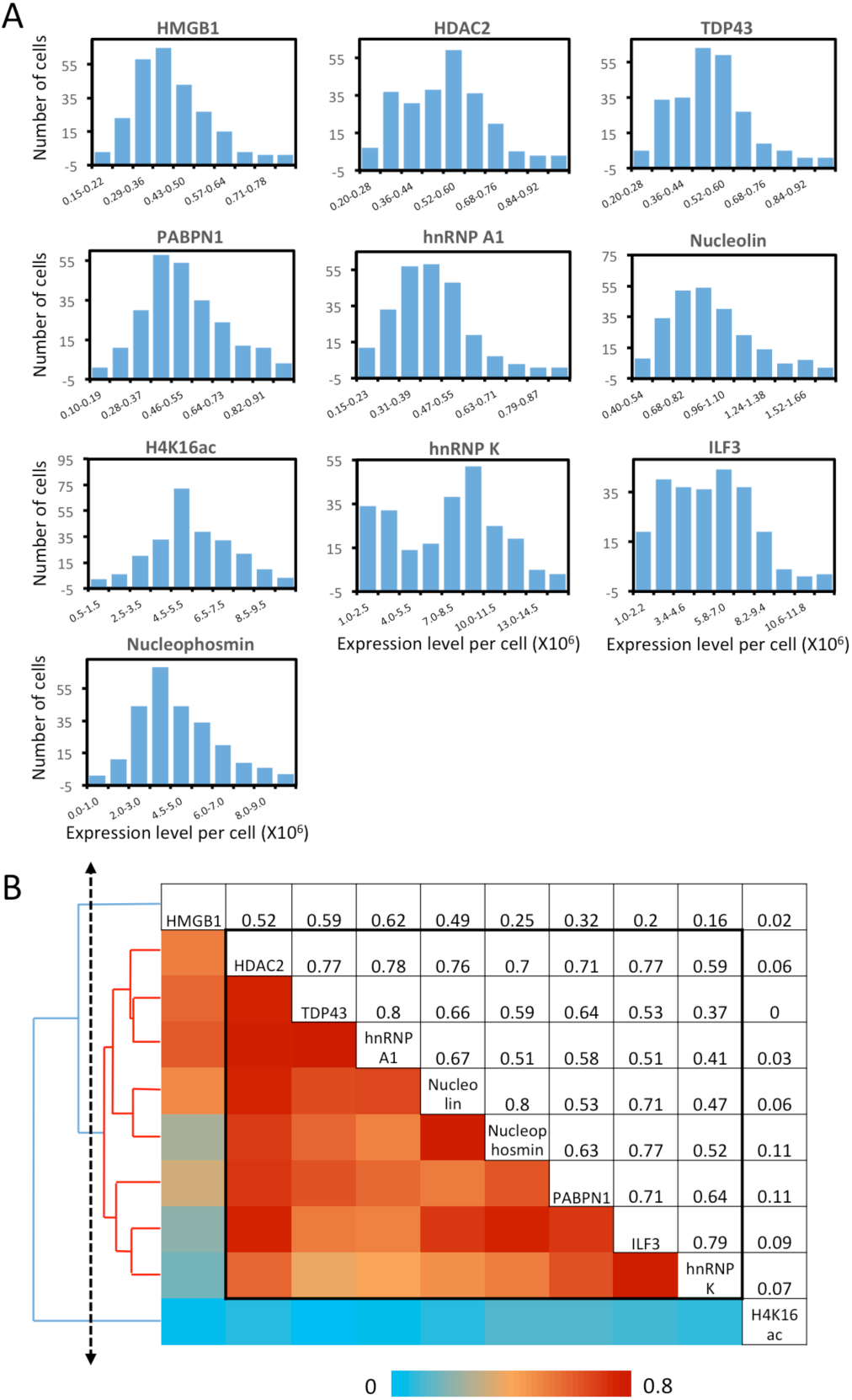
Protein expression heterogeneity and correlation. (A) Histograms of the expression level per cell of the 10 measured proteins. (B) Correlation of the expression levels of the 10 measured proteins and the hierarchical clustering tree. The upper triangle shows the expression correlation coefficient of each protein pair. The lower triangle displays the color corresponding to the correlation coefficient. And the protein names are shown in the diagonal. A group of proteins identified by a threshold on the cluster tree (dashed line) is indicated by the black box in the matrix and the red lines on the tree.

By analyzing expression covariation of different proteins, one can study which proteins are coregulated to elucidate their regulatory pathways. For bulk cell analysis, such studies usually require external stimuli to introduce varied gene expression among different cell populations. At the single-cell level, stochastic gene expression variation is generated in individual cells. By taking advantage of this natural expression fluctuation, one can perform single cell expression covariation analysis to refine existing regulatory pathways, suggest new regulatory pathways, and predict the function of unannotated proteins^38^. Appling this approach, we studied the pairwise expression correlation of the ten measured proteins (Supplementary Fig. 6), and calculated the correlation coefficient of each protein pair (Fig. 6B). Some of protein pairs show highly correlated covariation with correlation coefficients of ∼0.8, such as TDP43 and hnRNP A1 along with Nucleolin and Nucleophosmin. To further explore the regulatory network among the measured proteins, we adopted a hierarchical clustering approach^39^ (Fig. 6B). On the generated cluster tree, we identified a group of eight proteins with substantially correlated expression patterns (Fig. 6B). Indeed, all the eight proteins in this identified group are involved in the transcriptional regulation and processing related pathways^40–47^.

### Multiplexed in situ protein profiling in FFPE human hippocampus

The various cell types in the brain cooperate collectively to achieve high-order mental functions. To accurately observe and precisely manipulate brain activities, it is required to have much greater knowledge of the molecular identities of specific cell types. This knowledge is also fundamental to the discovery of the cell-type targeted therapy to treat brain disorders. The identities of neurons are determined by their locations, protein, RNA, and DNA profiles, etc. However, the current molecular classification of human neurons is only defined by single-cell RNA-seq^48,49^. No systematic analysis of neuronal heterogeneity has been reported based on protein expression or molecular profiling in their natural spatial contexts. Additionally, FFPE postmortem tissues are the major source of human brains with unlimited regional sampling and depth. However, the limited sensitivity of the existing multiplexed in situ protein analysis methods hinders their applications to profile the partially degraded proteins^50^ in highly autofluorescent FFPE tissues^26^.

To explore the human neuronal heterogeneity by multiplexed in suit protein profiling and also to assess the feasibility of applying CFT for analyzing FFPE tissues, we stained 8 proteins sequentially in the human hippocampus using HRP conjugated antibodies and tyramide-N_3_-Cy5. Of the 8 proteins, NeuN was selected as the neuronal marker51, and PABPN1, HMGB1, TDP43, hnRNP A1, hnRNP K, ILF3 along with Nucleophosmin were selected as the components of the transcriptional regulation and processing pathways^41,42,44–47,52^. Due to the high sensitivity of our approach, the imaging exposure time can be minimized without compromising the analysis accuracy. As a result, the whole tissue (∼1 cm × 1 cm) was imaged within 30 minutes in each cycle. With 8 reiterative staining cycles, all the proteins of interest were successfully detected in the tissue (Fig. 7). These results suggest that our approach enables multiplexed single-cell in situ protein profiling in FFPE tissues with high sample throughput and short assay time.

**Fig. 7.**
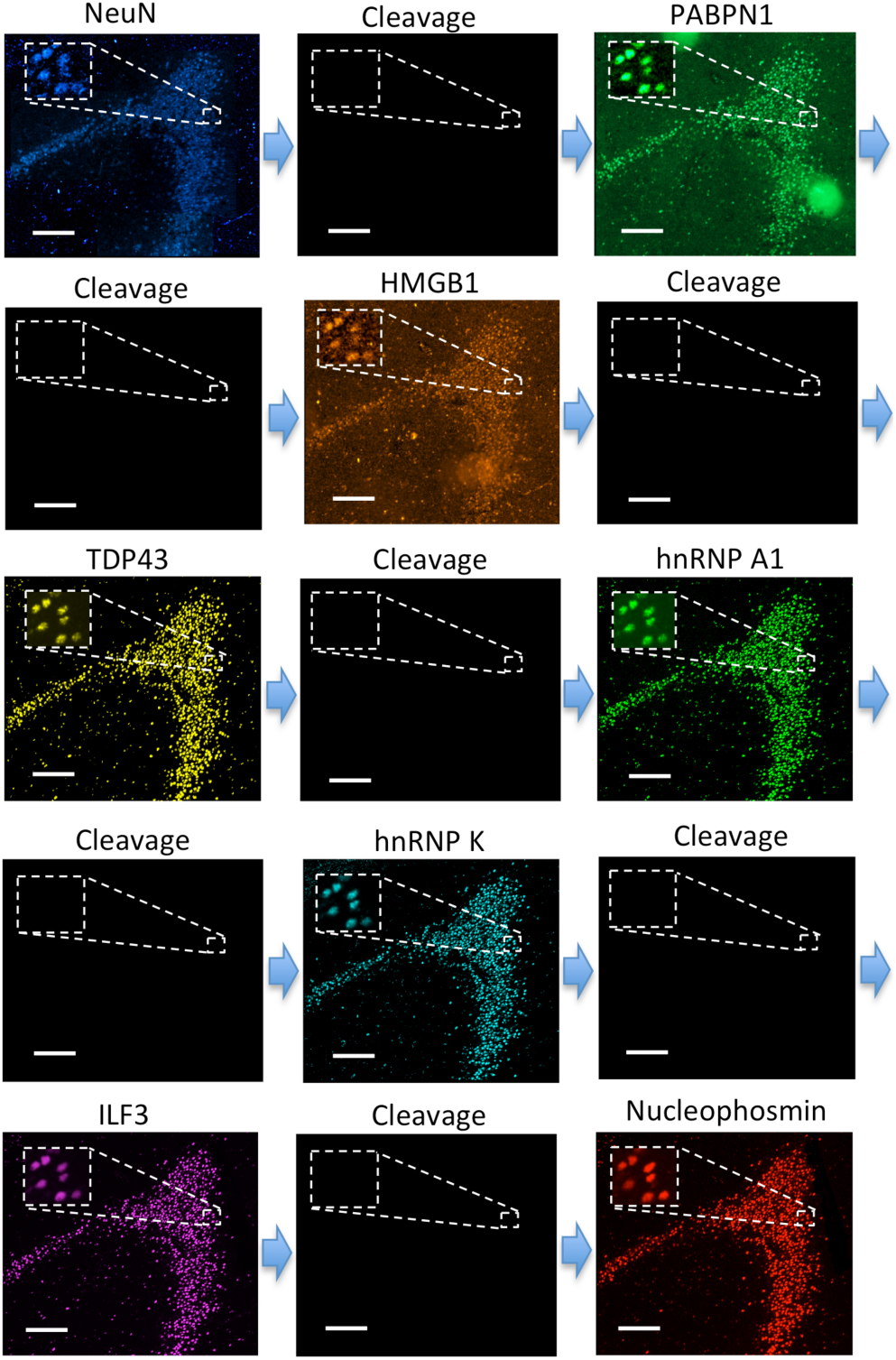
Eight different proteins are detected sequentially with HRP conjugated antibodies and tyramide-N3-Cy5 in the FFPE human brain tissue. Scale bars, 200 μm.

### Human neuronal heterogeneity and their spatial organization in the hippocampus

With the multiplexed single-cell in situ protein profiling results, we explored the neuronal heterogeneity and their spatial organization in the human hippocampus. In the examined tissue, we calculated the protein expression levels in >6000 individual neurons, which were identified by the neuronal marker NeuN^51^. We then applied the software viSNE^53^ to partition the individual neurons into 10 clusters (Fig. 8A) based on their protein expression profiles (Fig. 9 and Supplementary Fig. 7). By mapping these 10 clusters of cells back to their natural locations in the tissue (Fig. 8B, Supplementary Fig. 8,9), we observed that different subregions of the hippocampus consist of neurons from distinct clusters. For example, the dentate gyrus (DG) contains all the clusters except cluster 7, while the Cornu Ammonis (CA) fields are dominated by clusters 3, 6, 7, and 8. Within the CA fields, cluster 7 only appears in CA1, CA2 and CA3, but not in CA4 (Fig. 10A). In the DG, cluster 2 is the major cell class in the regions of interest (ROI) 1-5. In contrast, other subregions of the DG are mainly composed of clusters 1, 3, 4, 9 and 10 (Fig. 10B). These results suggest that our approach allows the investigation of the different cell type compositions and their spatial organizations in FFPE tissues.

**Fig. 8.**
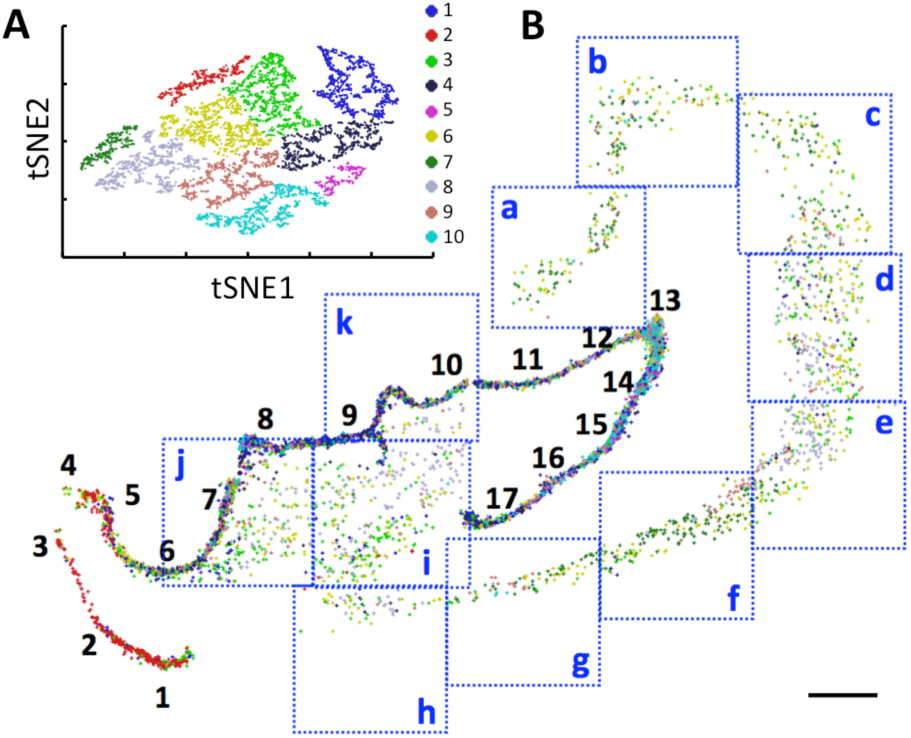
A) Over 6000 neurons in a human hippocampus are partitioned into 10 clusters. B) Anatomical locations of the individual neurons from the 10 clusters in the DG (1-17), CA1 (a-e), CA2 (f), CA3 (g,h) and CA4 (i-k). Scale bars, 2 mm.

**Fig. 9.**
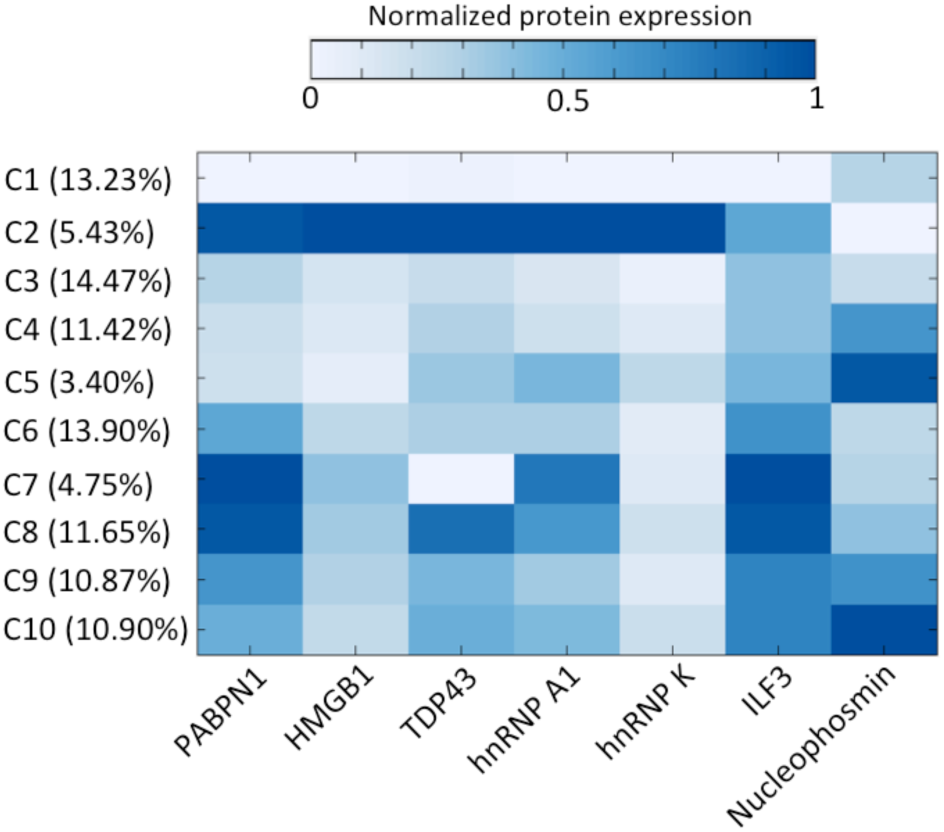
The distinct protein expression patterns in the 10 cell clusters and the percentage of cells in each cluster.

**Fig. 10.**
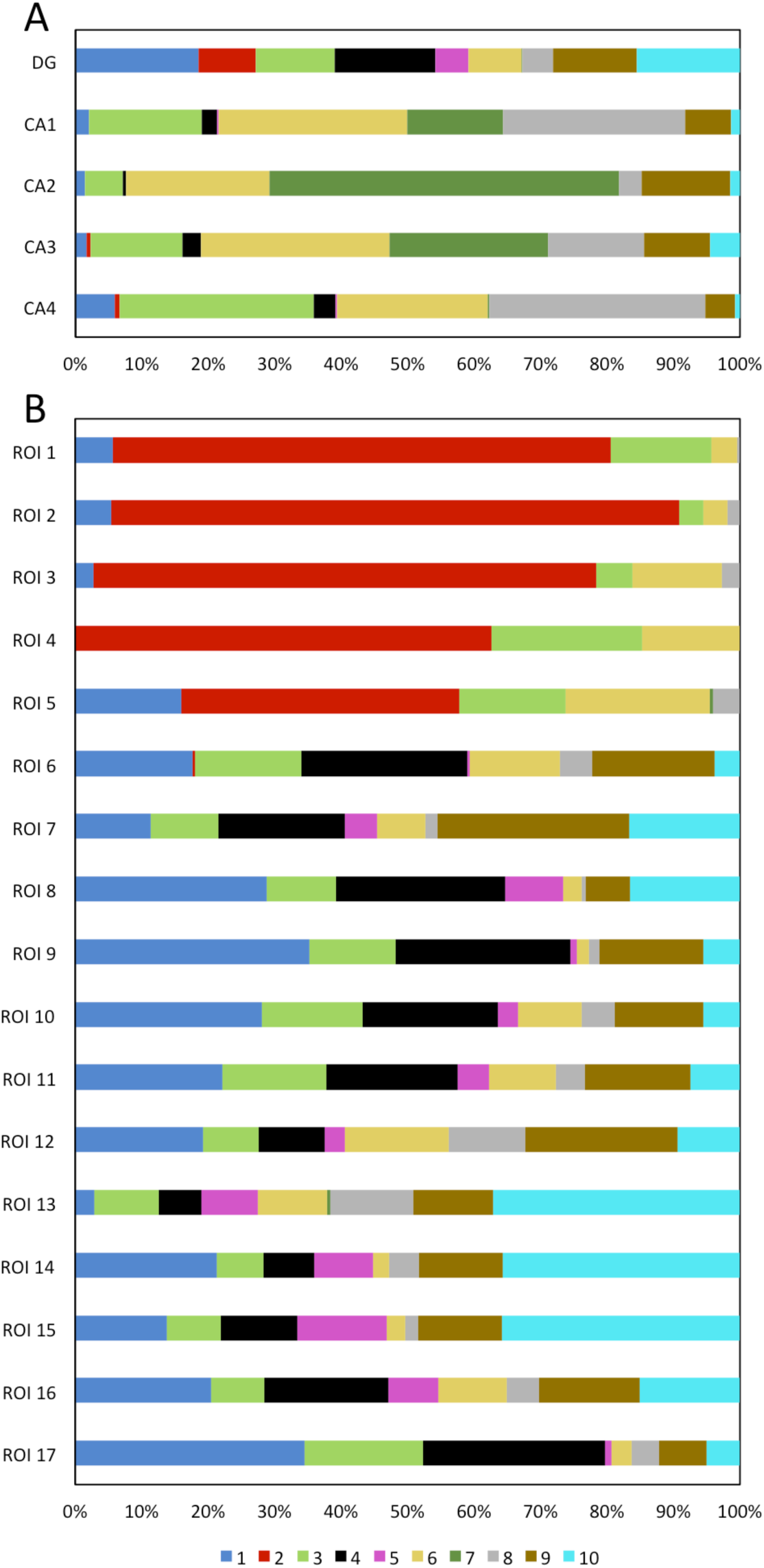
(A) The DG and CA fields are composed of neurons from different cell clusters. (B) Varied ROI in the DG are composed of neurons from different cell clusters.

## Discussion

In this study, we have designed and synthesized cleavable fluorescent tyramide, and applied it for multiplexed protein profiling in single cells of FFPE tissues in situ. Compared with the existing multiplexed protein imaging technologies, our approach has enhanced the detection sensitivity by 1-2 orders of magnitude. Additionally, by minimizing the imaging time and avoiding the pixel-by-pixel data acquisition, our method enables the whole-slide scanning within 30 minutes, which dramatically increases the sample throughput and reduces the assay time. Applying our approach, we have shown that different subregions of the human hippocampus consist of varied neuron clusters. Interestingly, these distinct clusters are defined only on the basis of the protein expression profiles, without incorporating the cellular spatial information into the clustering algorithm. These results suggest that the varied activity patterns and different microenvironment may contribute to the neuronal heterogeneity in the human hippocampus.

The multiplexing capacity of our in situ protein profiling approach depends on two factors: the cycling number and the number of proteins interrogated in each cycle. TCEP can efficiently remove the fluorophores within 30 minutes, while the antigenicity of protein targets is preserved after incubation with TCEP for more than 24 hours. These results suggest that at least ∼50 cycles can be carried out in one specimen. Coupled with the various established antibody stripping methods^54^ or HRP inactivation methods^55^, our approach will enable four or five different protein targets to be profiled in each analysis cycle using CFT with distinct fluorophores (Supplementary Fig. 10). Therefore, we envision that this CFT-based approach has the potential to detect hundreds of protein targets in the same tissue.

The cleavable fluorescent tyramide developed here can also be applied in other areas beyond protein analysis, such as DNA or RNA in situ hybridization^56^ and metabolic analysis^57^. The combination of these applications will enable the integrated DNA, RNA, protein and metabolic analysis at the optical resolution in intact tissues. Furthermore, coupled with a program-controlled microfluidic system^58^, a standard fluorescence microscope can be easily converted into an automatic highly multiplexed tissue imaging system. This comprehensive molecular imaging platform will bring new insights into cell signaling regulation, cell heterogeneity, cellular microenvironment, molecular diagnosis and cellular targeted therapy.

## Materials and Methods

### General information

Chemicals and solvents were purchased from Sigma-Aldrich or TCI America and were used directly without further purification, unless otherwise noted. Bioreagents were purchased from Invitrogen, unless otherwise indicated. 1H-NMR and 13C-NMR were taken on Varian Innova 500 MHz NMR spectrometers. Chemical shifts are reported in parts permillion (ppm) downfield from tetramethylsilane (TMS). Data are reported as follows: chemicalshift, multiplicity: singlet (s), doublet (d), triplet (t), multiplet (m), coupling constants *J* in Hz, and integration. HRMS was performed by the Arizona State University mass spectrometry facility.

### Synthesis of the tyramide-N3-Cy5

The compound 1 (Supplementary Fig. 11) prepared accordingly to the literature[21] was further purified by semi-preparative reverse phase HPLC [HPLC gradient: A, 100% 0.1 M TEAA; B, 100% MeCN; 0-2 min, 5% B (flow 2-5 ml/min); 2-15 min, 5-22% B (flow 5 ml/min); 15-20 min, 22-30% B (flow 5 ml/min); 20-30 min, 30-35% B (flow 5 ml/min); 30-32 min, 35-95% B (flow 5 ml/min); 32-35 min, 95% B (flow 5 ml/min); 35-37 min, 95-5% B (flow 5 ml/min); 37-40 min, 5% B (flow 5-2 ml/min)]. The fraction with retention time 25.6 min was collected and dried completely under reduced pressure. The purified compound 1 (3.9 mg, 4.86 μmol) was co-evaporated with anhydrous DMF (1 ml) and then dissolved in anhydrous DMF (300 μL). *N, N’*-disuccinimidyl carbonate (DSC) (6.2 mg, 24.3 μmol) in 40 μL of anhydrous DMF and 4-dimethylaminopyridine (DMAP) (3.0 mg, 24.3 μmol) were added to the above solution and the reaction mixture was stirred for 30 min at room temperature. Subsequently, to this reaction mixture tyramine hydrochloride (4.2 mg, 24.3 μmol) and *N,N*-diisopropylethylamine (DIPEA) (8.2 μL, 48.6 μmol) were added and the reaction mixture was stirred for 2 h at room temperature. After completion of the reaction, DMF was evaporated completely under reduced pressure. The crude product was purified by a preparative silica gel TLC plate (25 × 25 cm; silica gel 60; CH3OH:CH2Cl2 = 1:6; Rf = 0.2). Subsequently, the residue was dissolved in 0.1 M TEAA buffer/10% CH3CN followed by filtering off undissolved materials by nylon syringe filter (0.2 UM). Then the product was further purified by semi-preparative reverse phase HPLC [HPLC gradient: A, 100% 0.1 M TEAA; B 100% MeCN; 0-2 min, 5% B (flow 2-5 ml/min); 2-10 min, 5-22% B (flow 5 ml/min); 10-15 min, 22-30% B (flow 5 ml/min); 15-20 min, 30-40% B (flow 5 ml/min); 20-25 min, 40-50% B (flow 5 ml/min); 25-30 min, 50-60% B (flow 5 ml/min); 30-32 min, 60-70% B (flow 5 ml/min); 32-35 min, 70-95% B (flow 5 ml/min); 35-37 min, 95% B (flow 5 ml/min); 37-39 min, 95-5% B, (flow 5 ml/min); 39-42 min, 5% B (flow 5-2 ml/min)]. The fraction with retention time 14.1 min was collected and dried completely under reduced pressure. The residue was co-evaporated twice with water (2 ml) to afford compound 2 (1.1 mg, 24%) as a pure blue solid. ^1^H NMR (500 MHz, CD3OD) (Supplementary Fig. 12) δ 8.05-7.96 (m, 2H), 7.87-7.77 (m, 4H), 7.29 (dd, J = 22.1, 8.4 Hz, 2H), 6.98 (d, J = 7.4 Hz, 2H), 6.70 (d, J = 8.1 Hz, 2H), 6.55-6.47 (m, 1H), 6.22 (dd, J = 24.1, 13.4 Hz, 2H), 4.60 (t, J = 5.9 Hz, 1H), 4.16-3.97 (m, 6H), 3.89-3.84 (m, 1H), 3.73-3.64 (m, 2H), 3.21-3.10 (m, 4H), 2.59-2.52 (m, 2H), 2.19-2.12 (m, 2H), 1.83-1.70 (m, 4H), 1.70-1.53 (m, 12H), 1.35-1.22 (m, 6H); HRMS (ESI-, m/z) calcd for C47H58N7O11S2 [(M)^-^]: 960.3636, found: 960.3074.

### Cell culture

HeLa CCL-2 cells (ATCC) were maintained in Dulbelcco’s modified Eagle’s Medium (DMEM) supplemented with 10% fetal bovine serum, 100 U/mL penicillin and 100 g/mL streptomycin in a humidified atmosphere at 37 °C with 5% CO2. Cells were plated on chambered coverglass (0.2 ml medium/chamber) (Thermo Fisher Scientific) and allowed to reach 60% confluency in 1-2 days.

### Cell fixation

Cultured HeLa CCL-2 cells were fixed with 4% formaldehyde at 37°C for 15 min, permeabilized with 0.1% (vol/vol) Triton X-100 at room temperature for 15 min, and washed 3 times with 1X phosphate-buffered saline (PBS), each for 5 min.

### Endogenous peroxidase blocking

Fixed and permeabilized HeLa CCL-2 cells were incubated with 0.15% H2O2 in PBT (1X PBS, 0.1% (vol/vol) Triton X-100) for 10 min, and then washed 3 times with 1X PBS, each for 5 min.

### Immunofluorescence with CFT

Fixed and permeabilized HeLa CCL-2 cells were first blocked with 1X blocking buffer (1% (wt/vol) bovine serum albumin, 0.1% (vol/vol) Triton X-100, 10% (vol/vol) normal goat serum) at room temperature for 1 h. The cells were incubated with HRP conjugated primary antibodies at a concentration of 5 μg/mL in 1X blocking buffer for 45 min, and then washed 3 times with PBT, each for 5 min. Subsequently, cells were incubated with 10 pmol/μL tyramide-N3-Cy5 in amplification buffer (0.1 M Boric acid, pH=8.5) for 7 min. Cells were quickly washed twice with PBT, followed by 5 min wash with PBT for 3 times. Stained cells were washed with GLOX buffer (0.4% glucose and 10 mM Tris HCl in 2X saline-sodium citrate (SSC) buffer (300 mM sodium chloride, 30 mM trisodium citrate, pH = 7.0)) for 1 min at room temperature, and then imaged in GLOX solution (0.37 mg mL-1 glucose oxidase and 1% catalase in GLOX buffer). The used primary antibodies include HRP conjugated rabbit anti-HMGB1 (Thermo Fisher Scientific; PA5-22722), HRP conjugated rabbit anti-HDAC2 (Abcam; ab195851), HRP conjugated rabbit anti-TDP43 (Abcam; ab193850), HRP conjugated rabbit anti-PABPN1 (Abcam; ab207515), HRP conjugated rabbit anti-hnRNP A1 (Abcam; ab198535), HRP conjugated mouse anti-Nucleolin (Abcam; ab198492), HRP conjugated rabbit anti-Histone H4 (acetyl K16) (Abcam; ab200859), HRP conjugated mouse anti-hnRNP K (Abcam; ab204456), HRP conjugated rabbit anti-ILF3 (Abcam; ab206250) and HRP conjugated mouse anti-Nucleophosmin (Abcam; ab202579).

To stain protein Ki67, fixed and blocked HeLa CCL-2 cells were incubated with 5 μg/mL rabbit anti-Ki67 (Thermo Fisher Scientific; RB1510P1ABX) in 1X blocking buffer for 45 min, and then washed 3 times with PBT, each for 5 min. Afterwards, cells were incubated with 5 μg/mL HRP conjugated goat-anti-rabbit (Thermo Fisher Scientific; A16110) in 1% (wt/vol) bovine serum albumin in PBT for 30 min, followed by 3 times wash with PBT, each for 5 min. Subsequently, cells were incubated with 10 pmol/μL tyramide-N3-Cy5 in amplification buffer for 7 min. Cells were quickly washed twice with PBT, followed by 5 min wash with PBT for 3 times. Cells were then imaged in GLOX solution.

### Fluorophore cleavage and HRP deactivation

To remove the fluorophores and simultaneously deactivate horseradish peroxidase (HRP), cells were incubated with tris(2-carboxyethyl)phosphine (TCEP) (100 mM, pH=9.5) at 50°C for 30 minutes. To explore the cleavage efficiencies under different temperatures, cells were incubated with TCEP (100 mM, pH=9.5) at 37°C, 50°C and 65°C for 30 minutes. To study the cleavage kinetics, cells were incubated with TCEP (100 mM, pH=9.5) at 50°C for 5, 15, 30 and 60 minutes. Following the TCEP incubation, cells were washed 3 times with PBT and 3 times with 1X PBS, each for 5 min. Cells were then imaged in GLOX solution. To evaluate the HRP deactivation efficiencies following the TCEP incubation, cells were incubated with 10 pmol/μL tyramide-N3-Cy5 in amplification buffer for 7 min. After 2 times quick wash and 3 times 5 min wash with PBT, cells were imaged in GLOX solution.

### Conventional immunofluorescence

The Cy5 labeled primary and secondary antibodies were prepared accordingly to the literature^[1]^. For direct immunofluorescence, fixed and blocked HeLa CCL-2 cells were incubated with 5 μg/mL Cy5 labeled rabbit anti-Ki67 primary antibodies (Thermo Fisher Scientific; RB1510P1ABX) in the 1X blocking buffer for 45 min at room temperature. Cells were washed 3 times with PBT, each for 5 min, and then imaged. For indirect immunofluorescence, fixed and blocked HeLa CCL-2 cells were incubated with 5 μg/mL rabbit anti-Ki67 (Thermo Fisher Scientific; RB1510P1ABX) for 45 min in 1X blocking buffer, then washed 3 times with PBT, each for 5 min. Then cells were incubated with 5 μg/mL Cy5 labeled goat-anti-rabbit (Thermo Fisher Scientific; A16112) in 1% (wt/vol) bovine serum albumin in PBT for 30 min, followed by 3 times wash with PBT, each for 5 min. Cells were then imaged in GLOX solution.

### Multiplexed protein analysis with CFT in cells

Fixed and blocked HeLa CCL-2 cells were incubated with 5 μg/mL HRP conjugated primary antibodies at room temperature for 45 min, and then stained by tyramide-N3-Cy5. After imaging, stained cells were incubated with 100 mM TCEP (pH=9.5) at 50°C for 30 min, followed by the next immunofluorescence cycle. The sequentially used primary antibodies include HRP conjugated rabbit anti-HMGB1 (Thermo Fisher Scientific; PA5-22722), HRP conjugated rabbit anti-HDAC2 (Abcam; ab195851), HRP conjugated rabbit anti-TDP43 (Abcam; ab193850), HRP conjugated rabbit anti-PABPN1 (Abcam; ab207515), HRP conjugated rabbit anti-hnRNP A1 (Abcam; ab198535), HRP conjugated mouse anti-Nucleolin (Abcam; ab198492), HRP conjugated rabbit anti-Histone H4 (acetyl K16) (Abcam; ab200859), HRP conjugated mouse anti-hnRNP K (Abcam; ab204456), HRP conjugated rabbit anti-ILF3 (Abcam; ab206250) and HRP conjugated mouse anti-Nucleophosmin (Abcam; ab202579). For control experiments, fixed and blocked HeLa CCL-2 cells were incubated with 5 μg/mL HRP conjugated primary antibodies at room temperature for 45 min, and then stained by Cy5 labeled tyramide (PerkinElmer).

### Deparaffinization and antigen retrieval

The brain formalin-fixed paraffin-embedded (FFPE) tissue slide was deparaffinized 3 times in xylene, each for 10 min. Then, the slide was immersed in 100% ethanol for 2 min, 95% ethanol for 1 min, 70% ethanol for 1 min, 50% ethanol for 1 min, 30% ethanol for 1 min, and rinsed with deionized water. Subsequently, the slide was immersed in antigen retrieval buffer (10 mM sodium citrate, 0.05% Tween 20, pH=6.0), and water-bathed in a pressure cooker for 20 min with the “high pressure” setting. Afterwards, the slide was rinsed 3 times with 1X PBS, each for 5 min.

### Multiplexed protein staining in FFPE tissues

After deparaffinization and antigen retrieval, the brain FFPE tissue was first blocked by 0.15% H2O2 for 10 min and then washed 3 times with 1X PBS, each for 5 min. The tissue was then blocked in 1X blocking buffer at room temperature for 1 h. Subsequently, the tissue was incubated with 5 μg/mL biotin conjugated Rabbit anti-NeuN (Abcam; ab204681) in 1X blocking buffer for 45 min, and then washed 3 times with PBT, each for 5 min. Afterwards, the tissue was incubated with 5 μg/mL HRP conjugated streptavidin (Abcam; ab7403) in 1% (wt/vol) bovine serum albumin in PBT for 30 min, followed by 3 times wash with PBT, each for 5 min. Subsequently, the tissue was incubated with 10 pmol/μL tyramide-N3-Cy5 in amplification buffer for 7 min. The tissue was quickly washed twice with PBT, followed by 5 min wash with PBT for 3 times. After imaging, the tissue was incubated with 100 mM TCEP (pH=9.5) at 50°C for 30 min. The tissue was imaged again before initiating the next cycle. In the following cycles, the tissue was incubated with 5 μg/mL HRP conjugated primary antibodies in 1X blocking buffer for 45 min. Subsequently, the tissue was stained with tyramide-N3-Cy5 and imaged. After incubated with 100 mM TCEP (pH=9.5) at 50°C for 30 min, the tissue was imaged again, followed by the next analysis cycle. The sequentially used primary antibodies include HRP conjugated rabbit anti-PABPN1 (Abcam; ab207515), HRP conjugated rabbit anti-HMGB1 (Thermo Fisher Scientific; PA5-22722), HRP conjugated rabbit anti-TDP43 (Abcam; ab193850), HRP conjugated rabbit anti-hnRNP A1 (Abcam; ab198535), HRP conjugated mouse anti-hnRNP K (Abcam; ab204456), HRP conjugated rabbit anti-ILF3 (Abcam; ab206250) and HRP conjugated mouse anti-Nucleophosmin (Abcam; ab202579).

### Imaging and data analysis

Both stained cells and the FFPE brain tissue were imaged under a Nikon Ti-E epifluorescence microscope equipped with 20X objective. Images were taken using a CoolSNAP HQ2 camera and Chroma filter 49009. Cell segmentation and intensity quantification were processed by NIS-Elements Imaging software. Pseudo-color images were generated using ImageJ. Protein expression heterogeneity and correlation were analyzed with Excel (Microsoft). The hierarchical clustering was performed with Cluster 3.0 (http://bonsai.hgc.jp/~mdehoon/software/cluster/). ViSNE maps were generated from CYT (https://www.c2b2.columbia.edu/danapeerlab/html/cyt.html).[53]

## Acknowledgments

This work is supported by the National Institute of General Medical Sciences (1R01GM127633), the National Institute Of Allergy And Infectious Diseases (R21AI132840), Arizona State University startup funds, and Arizona State University/Mayo Clinic seed grant (ARI-219693).

