## Supplemental Information for "Highly sensitive in situ proteomics with cleavable fluorescent tyramide reveals human neuronal heterogeneity"

.

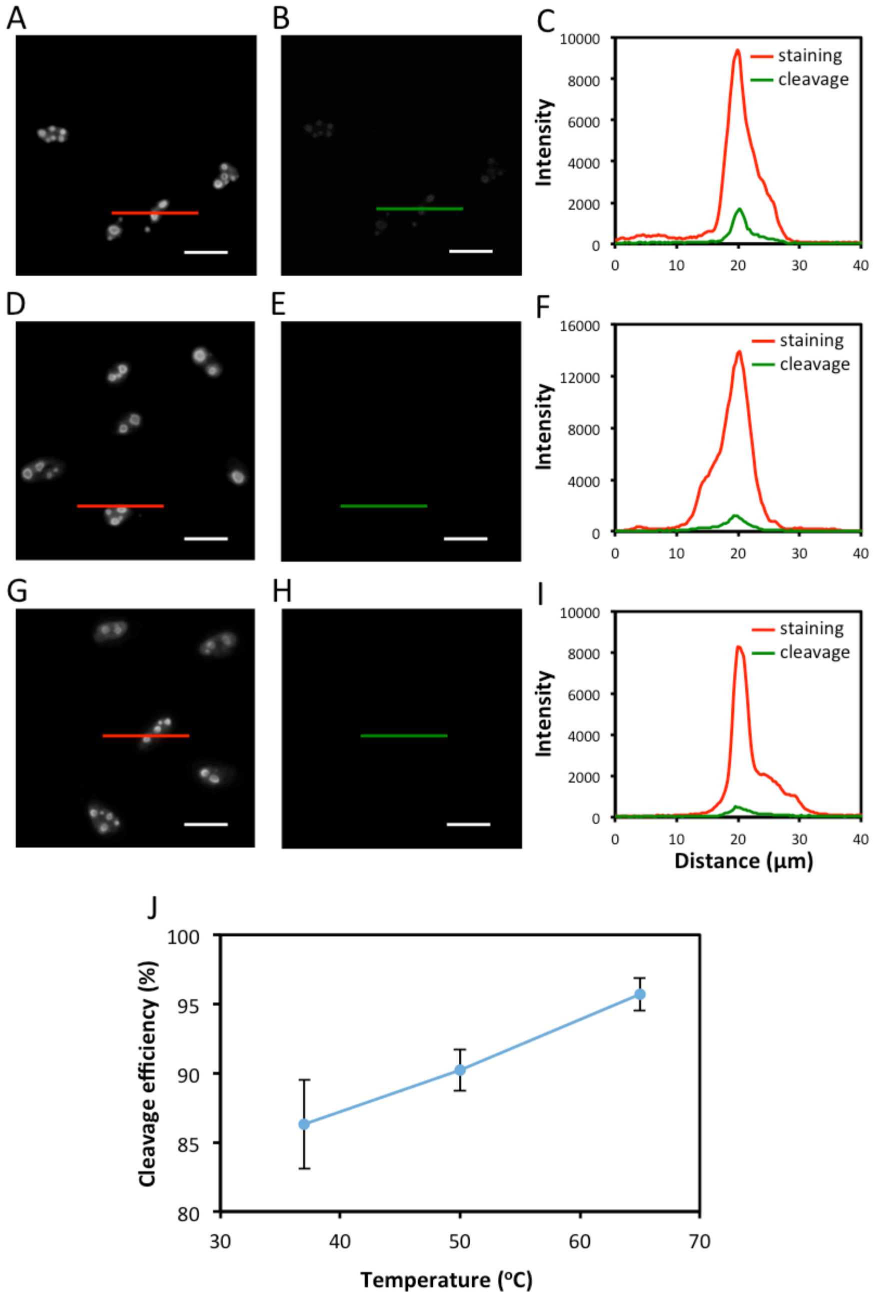


**Supplementary Figure 1**. (A) Protein Ki67 in HeLa cells is stained with tyramide-$N_3$-Cy5. (B) The stained cells are incubated with TCEP at 37°C for 30 minutes. (C) Fluorescence intensity profile corresponding to the red line and green line positions in (A) and (B). (D) Protein Ki67 in HeLa cells is stained with tyramide-$N_3$-Cy5. (E) The stained cells are incubated with TCEP at 50°C for 30 minutes. (F) Fluorescence intensity profile corresponding to the red line and green line positions in (D) and (E). (G) Protein Ki67 in HeLa cells is stained with tyramide-$N_3$-Cy5. (H) The stained cells are incubated with TCEP at 65°C for 30 minutes. (I) Fluorescence intensity profile corresponding to the red line and green line positions in (G) and (H). (J) Fluorophore cleavage efficiency at different reaction temperatures (n = 30 positions). Scale bars, 20 µm.

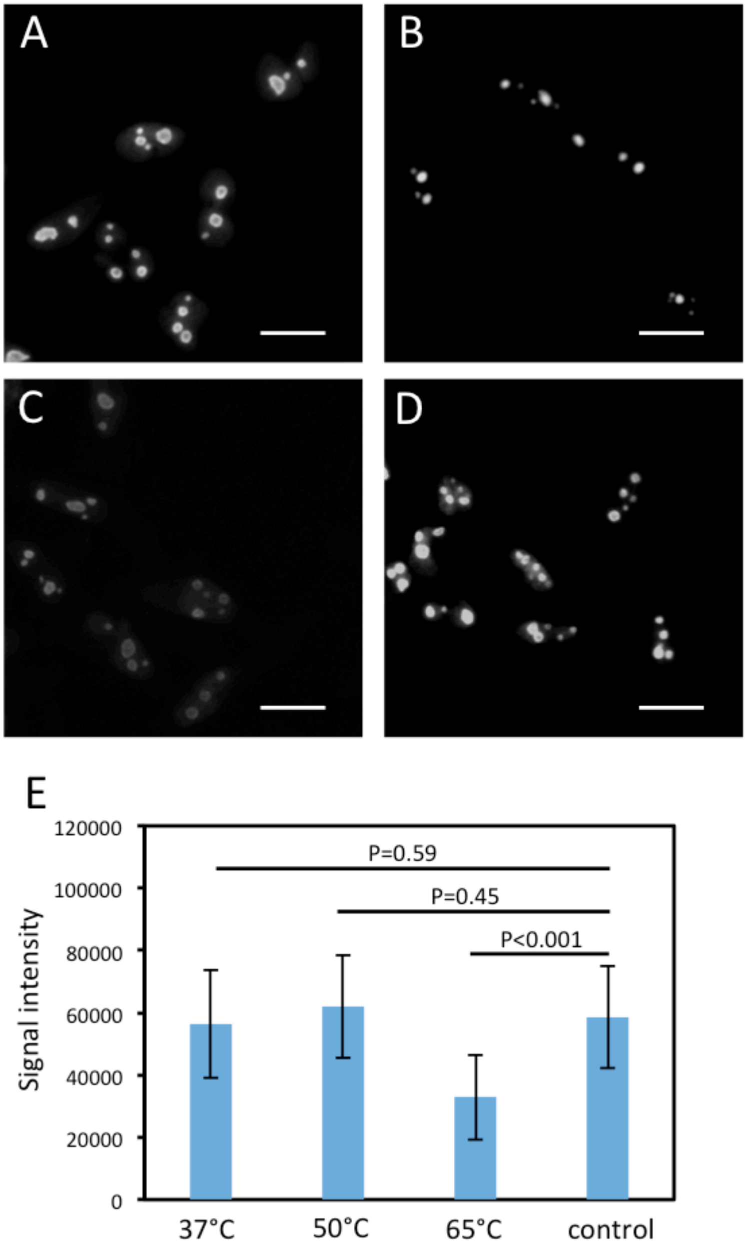


**Supplementary Figure 2**. After incubation with TCEP at (A) 37°C, (B) 50°C and (C) 65°C for 24 hours, or (D) without any TCEP pre-treatment, protein Ki67 in HeLa cells is stained with tyramide-$N_3$-Cy5. (E) The obtained signal intensities with TCEP pre-treatment at different temperatures and without any pre-treatment (control) (n = 30 positions). Scale bars, 20 µm.

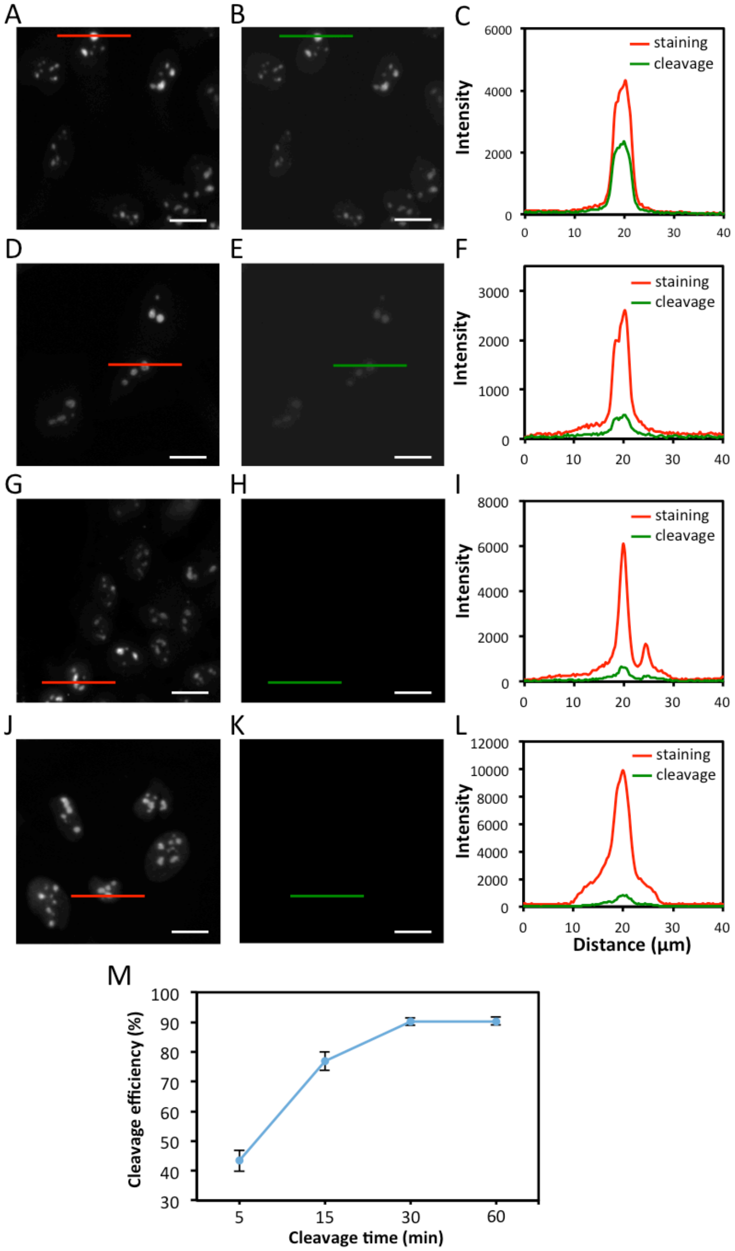


**Supplementary Figure 3**. A) Protein Ki67 in HeLa cells is stained with tyramide-$N_3$-Cy5. (B) The stained cells are incubated with TCEP at 50°C for 5 minutes. (C) Fluorescence intensity profile corresponding to the red line and green line positions in (A) and (B). (D) Protein Ki67 in HeLa cells is stained with tyramide-$N_3$-Cy5. (E) The stained cells are incubated with TCEP at 50°C for 15 minutes. (F) Fluorescence intensity profile corresponding to the red line and green line positions in (D) and (E). (G) Protein Ki67 in HeLa cells is stained with tyramide-$N_3$-Cy5. (H) The stained cells are incubated with TCEP at 50°C for 30 minutes. (I) Fluorescence intensity profile corresponding to the red line and green line positions in (G) and (H). (J) Protein Ki67 in HeLa cells is stained with tyramide-$N_3$-Cy5. (K) The stained cells are incubated with TCEP at 50°C for 60 minutes. (L) Fluorescence intensity profile corresponding to the red line and green line positions in (J) and (K). (M) Fluorophore cleavage efficiency at different reaction time (n = 30 positions). Scale bars, 20 µm.

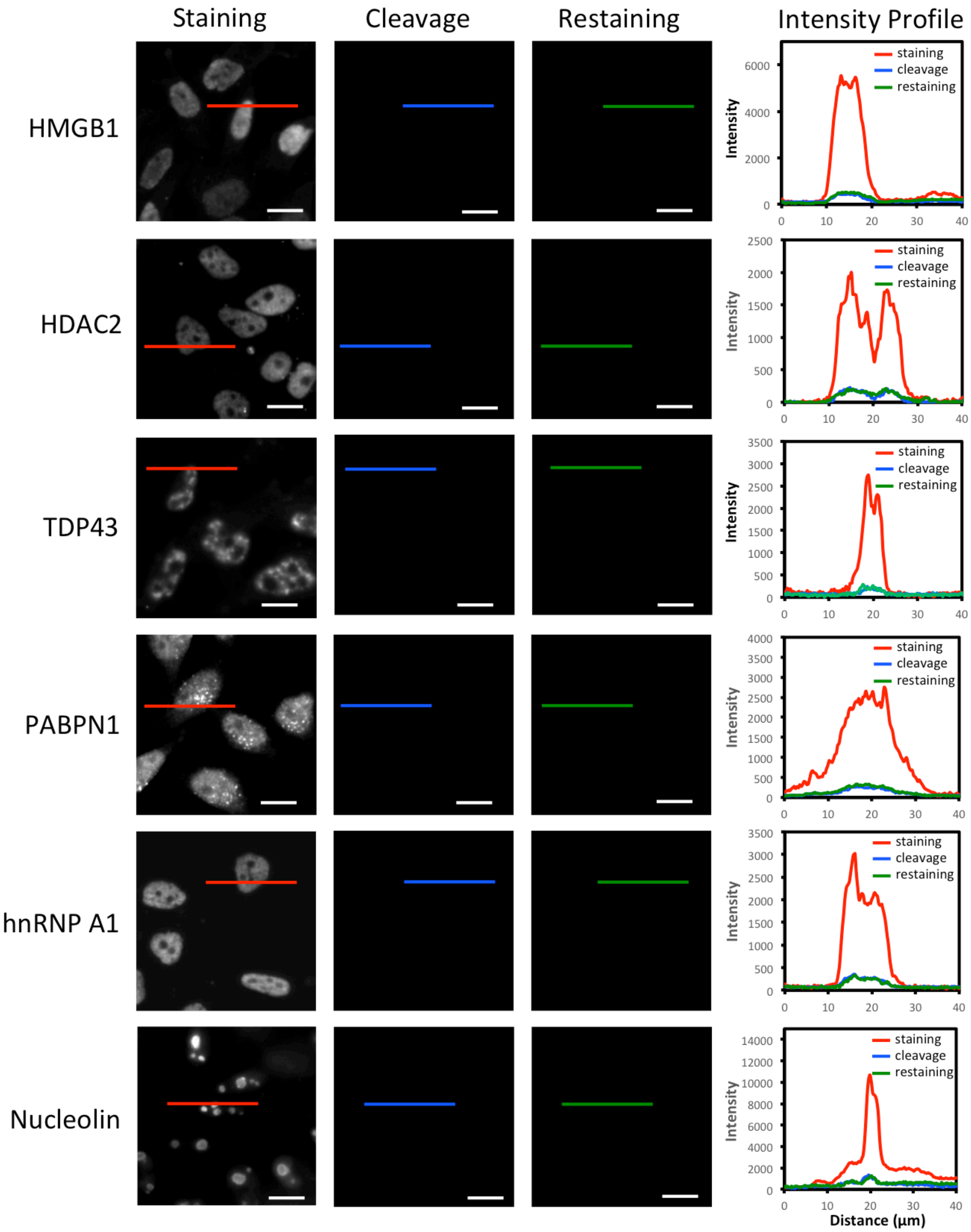

Staining
Cleavage
Restaining
Intensity Profile
HMGB1
HDAC2
TDP43
PABPN1
hnRNP A1
Nucleolin
staining
cleavage
restaining
Intensity
Distance (μm)

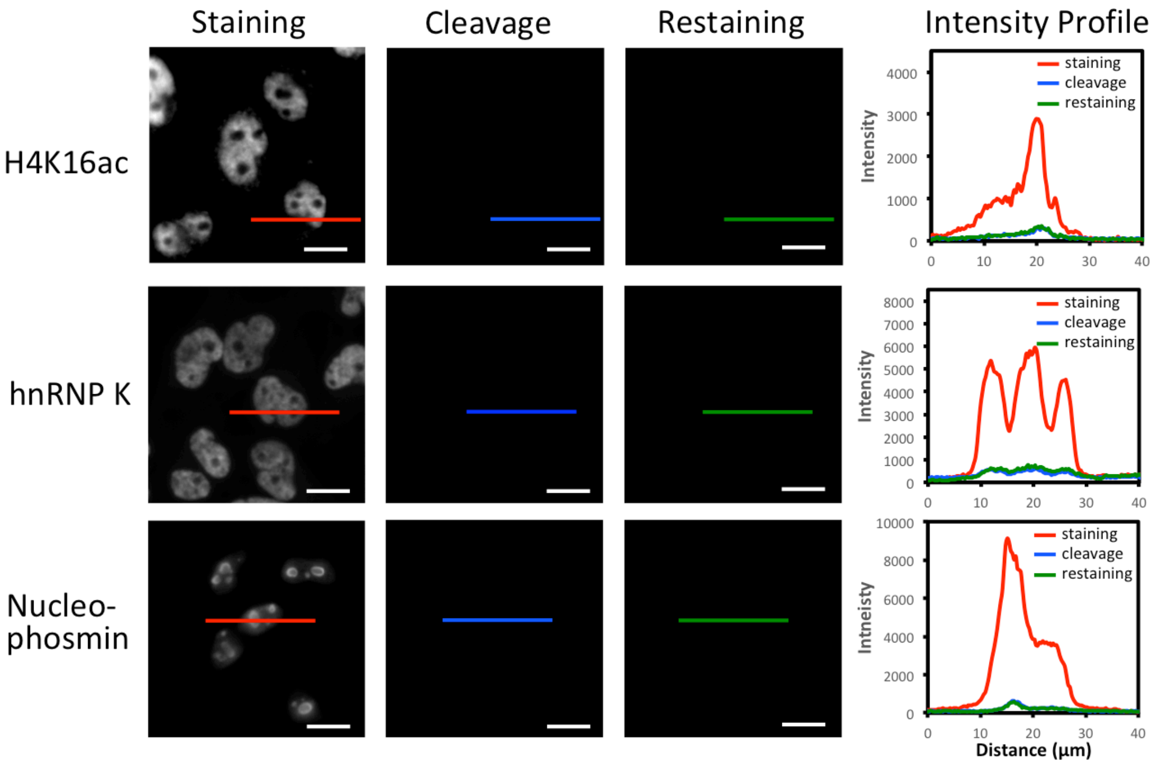


**Supplementary Figure 4**. Different proteins in HeLa cells are stained with HRP conjugated antibodies and tyramide-$N_3$-Cy5 (the first column). The stained cells are incubated with TCEP (the second column). Subsequently, the cells are incubated with tyramide-$N_3$-Cy5, again (the third column). Fluorescence intensity profiles corresponding to the red, blue and green line positions in the staining, cleavage and restaining images (the fourth column). Scale bars, 15 µm.

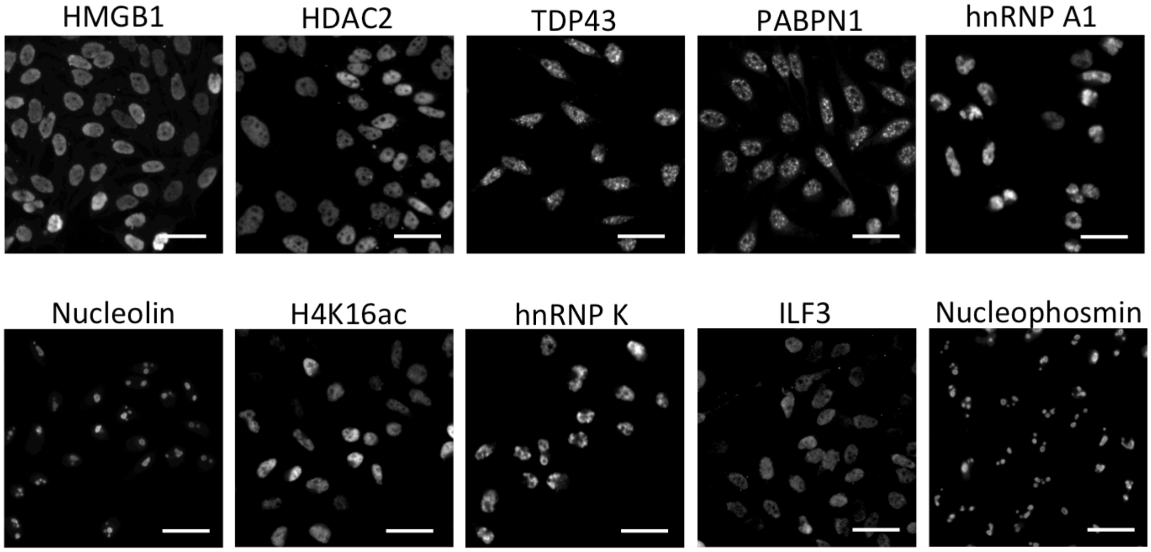


**Supplementary Figure 5**. 10 different proteins are stained with the corresponding HRP conjugated antibodies and Cy5 labeled tyramide in different HeLa cells. Scale bars, 40 μm.

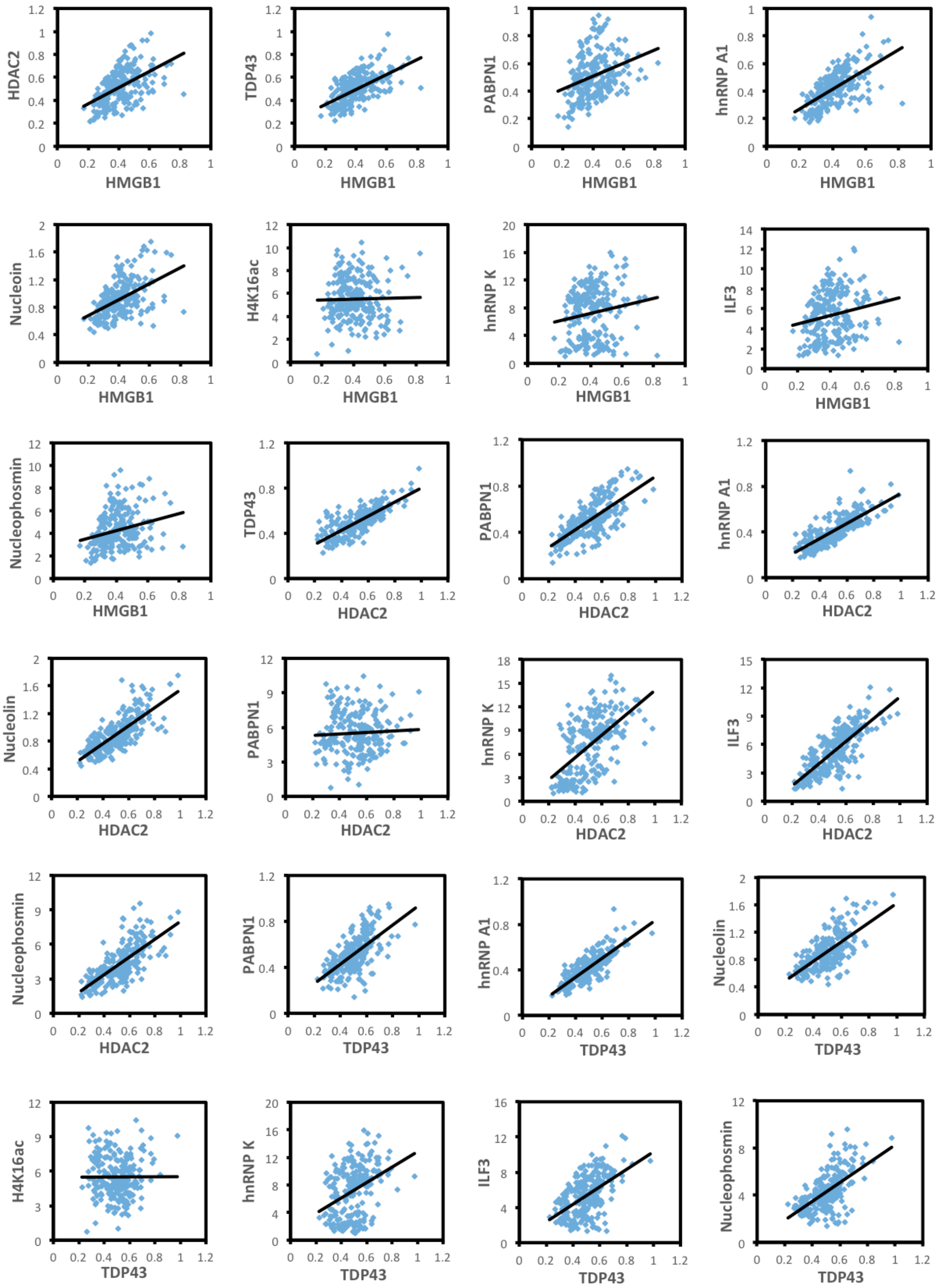
HDAC2
HMGB1
TDP43
HMGB1
PABPN1
HMGB1
hnRNP A1
HMGB1
Nucleoin
HMGB1
H4K16ac
HMGB1
hnRNP K
HMGB1
ILF3
HMGB1
Nucleophosmin
HMGB1
TDP43
HDAC2
PABPN1
HDAC2
hnRNP A1
HDAC2
Nucleolin
HDAC2
PABPN1
HDAC2
hnRNP K
HDAC2
ILF3
HDAC2
Nucleophosmin
HDAC2
PABPN1
TDP43
hnRNP A1
TDP43
Nucleolin
TDP43
H4K16ac
TDP43
hnRNP K
TDP43
ILF3
TDP43
Nucleophosmin
TDP43

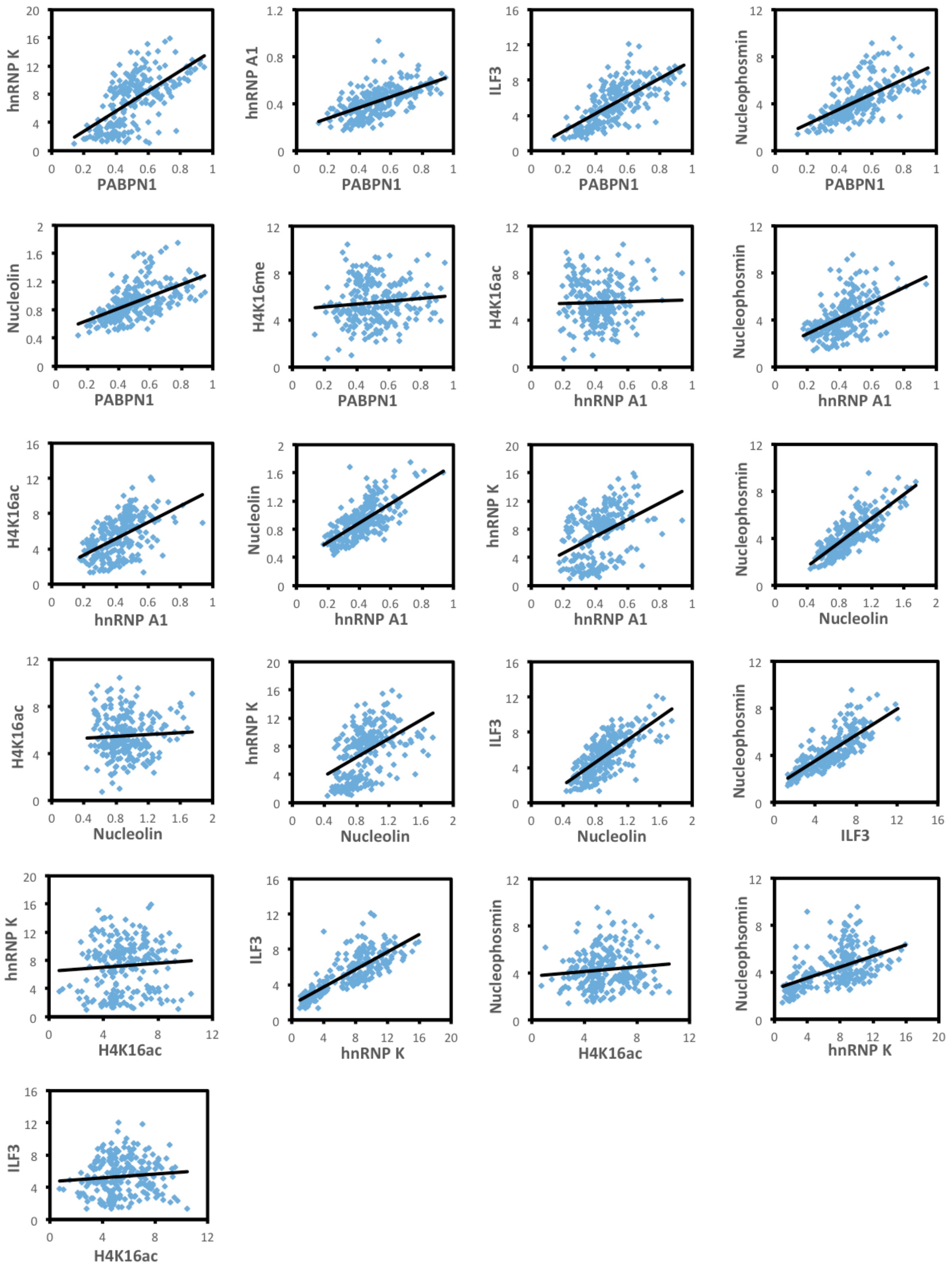


**Supplementary Figure 6**. Raw expression correlation data of each gene pair. Each spot corresponds to one cell with expression levels in the x and y axes ($\times 10^6$).

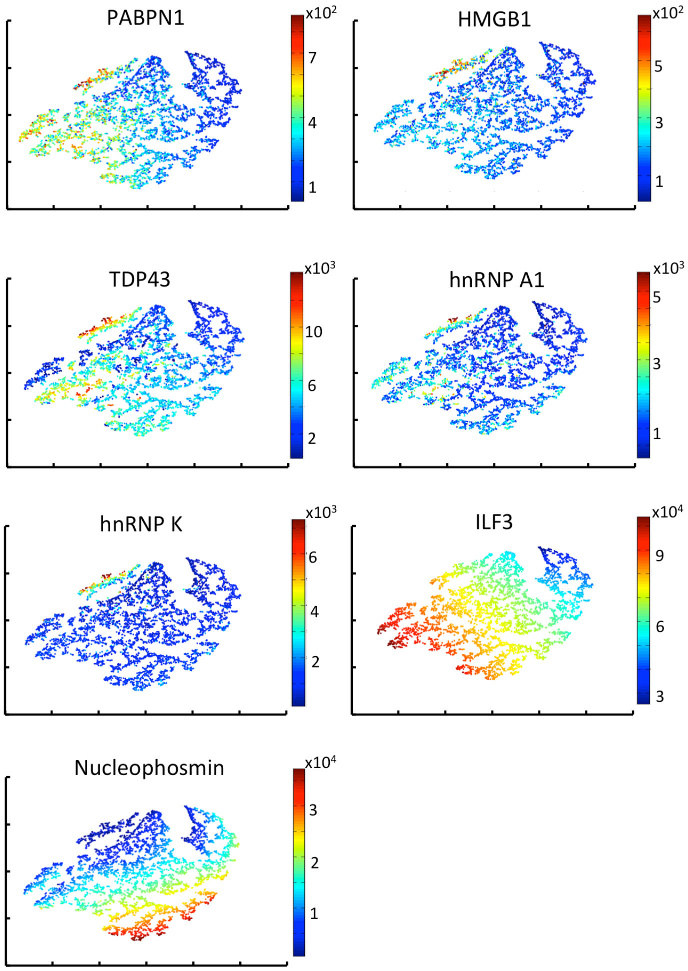


**Supplementary Figure 7**. Distribution of single-cell protein expression in viSNE plots.

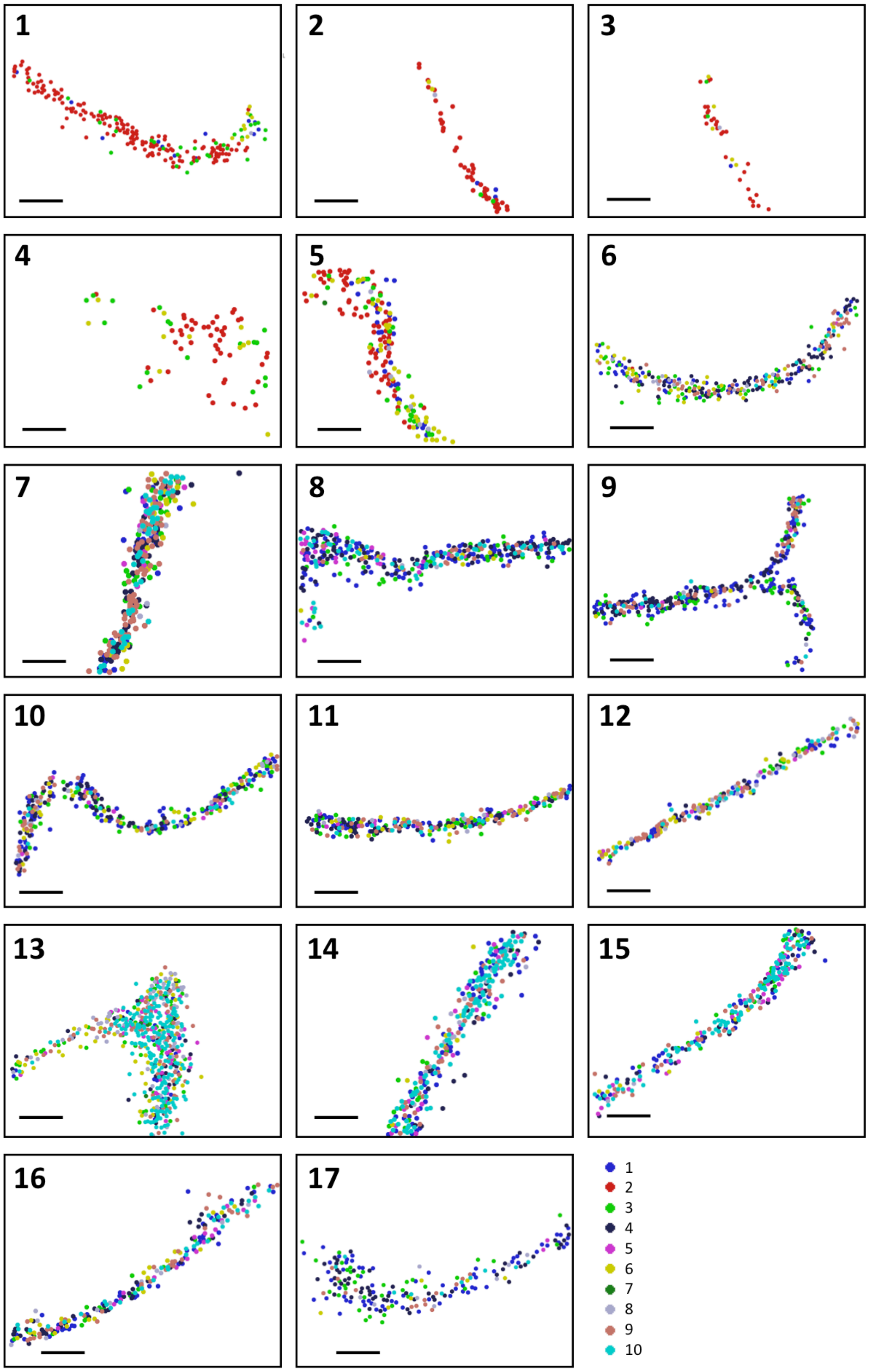


**Figure S8**. Zoom-in views of different regions of interest (ROI) in the dentate gyrus (DG) in Figure 8B. Scale bars, 200 μm.

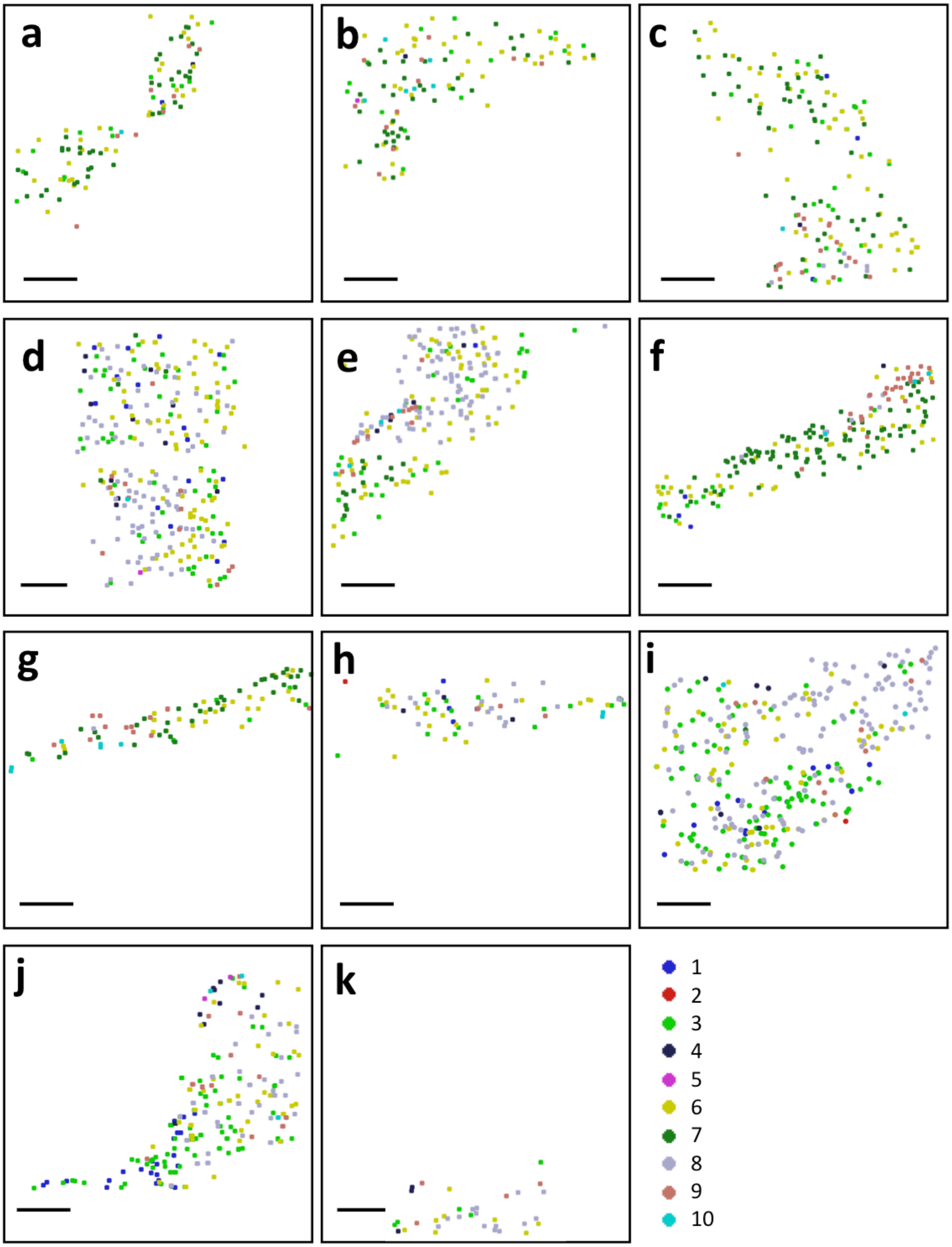


**Supplementary Figure 9**. Zoom-in views of different ROI in the Cornu Ammonis (CA) fields in Figure 8B. Scale bars, 500 µm.

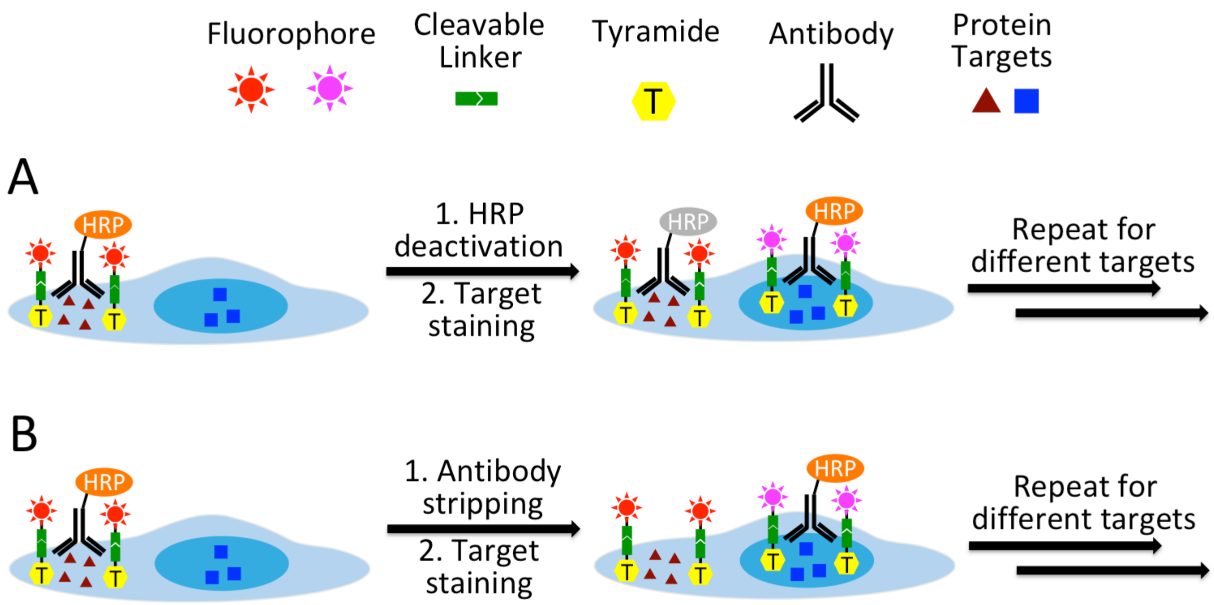


**Supplementary Figure 10**. (**A**) Through reiterative HRP deactivation or (**B**) cyclic antibody stripping, multiple protein targets can be detected in each analysis cycle using CFT with different fluorophores.

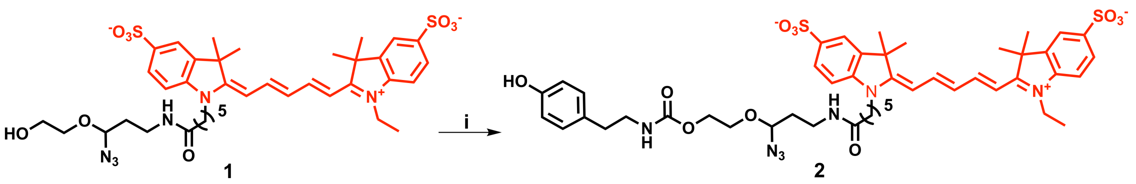


**Supplementary Figure 11**. Synthesis of tyramide-$N_3$-Cy5. Reagents and conditions: (i) DSC, DMAP, DMF, rt, 30 min; and then tyramine hydrochloride, DIPEA, rt, 2 h.

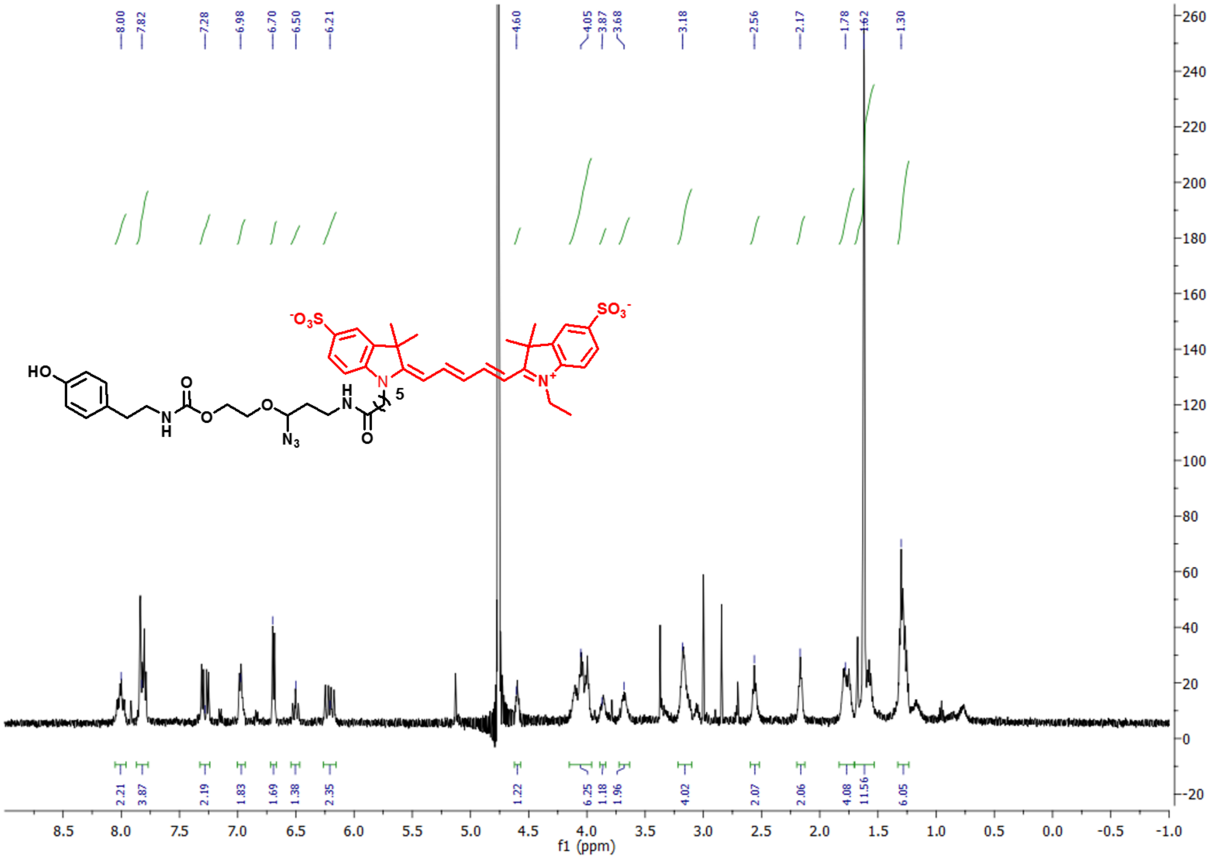

**Supplementary Figure 12**. $^{1}$H NMR of tyramide-N3-Cy5 (500 MHz, $CD_3OD$)
